# Next-generation large-scale binary protein interaction network for *Drosophila*

**DOI:** 10.1101/2022.08.02.502359

**Authors:** Hong-Wen Tang, Kerstin Spirohn, Yanhui Hu, Tong Hao, István A. Kovács, Yue Gao, Richard Binari, Donghui Yang-Zhou, Kenneth H. Wan, Joel S. Bader, Dawit Balcha, Wenting Bian, Benjamin W. Booth, Atina G. Cote, Steffi De Rouck, Alice Desbuleux, Dae-Kyum Kim, Jennifer J. Knapp, Wen Xing Lee, Irma Lemmens, Cathleen Li, Mian Li, Roujia Li, Hyobin Julianne Lim, Katja Luck, Dylan Markey, Carl Pollis, Sudharshan Rangarajan, Jonathan Rodiger, Sadie Schlabach, Yun Shen, Bridget TeeKing, Frederick P. Roth, Jan Tavernier, Michael A. Calderwood, David E. Hill, Susan E. Celniker, Marc Vidal, Norbert Perrimon, Stephanie E. Mohr

## Abstract

Generating reference maps of the interactome networks underlying most cellular functions can greatly illuminate genetic studies by providing a protein-centric approach to finding new components of existing pathways, complexes, and processes. Here, we applied state-of-the-art experimental and bioinformatics methods to identify high-confidence binary protein-protein interactions (PPIs) for *Drosophila melanogaster*. We performed four all-by-all yeast two-hybrid (Y2H) screens of >10,000 *Drosophila* proteins, resulting in the ‘FlyBi’ dataset of 8,723 PPIs among 2,939 proteins. As part of this effort, we tested subsets of our data and data from previous PPI datasets using an orthogonal assay, which allowed us to normalize data quality across datasets. Next, we integrated our FlyBi data with previous PPI data, resulting in an expanded, high-confidence binary *Drosophila* reference interaction network, DroRI, comprising 17,232 interactions among 6,511 proteins. These data are accessible through the Molecular Interaction Search Tool (MIST) and other databases. To assess the utility of the PPI resource, we used novel interactions from the FlyBi dataset to generate an autophagy interaction network that we validated *in vivo* using two different autophagy-related assays. We found that *deformed wings* (*dwg*) encodes a protein that is both a regulator and a target of autophagy. Altogether, the resources generated in this project provide a strong foundation for building high-confidence new hypotheses regarding protein networks and function.

## Main Text

Protein-protein interactions (PPIs) are central to cell biological processes, such as formation of multiprotein complexes and enzymes, receptor-ligand and kinase-substrate interactions, intracellular signal transduction, and regulation of transcription and translation. A number of complementary methods can be used to identify PPIs, including mass spectrometry-based methods for identification of protein complexes and two-component methods such as yeast two-hybrid (Y2H) analysis for identification of binary interactions^1^. Results from systematically screened and validated binary interactions contribute to the development of specific hypotheses regarding the functional *in vivo* relevance of individual PPIs. Moreover, when applied at large scale and integrated with other datasets, networks of binary interactions elucidate new components of known pathways. Particularly relevant to this study, since the release of the last binary PPI map for *Drosophila melanogaster* two decades ago^2^, methods for identification of binary interactions have improved and caveats to the approach are now well understood^3^. Innovations in experimental approach and analysis, as well as production of proteome-scale open reading frame (ORF) clone collections, made it possible to increase both the scale and the quality of binary interaction screens. Indeed, simply increasing the number of ORFs tested in Y2H assays contributes to new discoveries and brings protein-centric studies closer to the scale that can be accomplished with nucleic acid-based studies such as transcriptomics analyses. In recognition of the value of binary protein information to research study, binary interaction methods have been applied at an increasingly large scale for the discovery of PPI networks for several proteomes, including the human and yeast proteomes^4, 5^.

*Drosophila melanogaster* is an exemplary research system with a rich history of impactful contributions to our understanding of conserved biological processes and enduring relevance in biological and biomedical research^6–8^. The *Drosophila* research community has made significant investments in technology and resource development in addition to research studies, leading to a wealth of available genetic methods, fly stock reagents, large-scale datasets, and databases that can be used as research tools and mined for new hypothesis development, disease modeling, and experimental studies^9^. These include genome-wide genetic and RNAi screens^10–12^; extensive genomics studies^13, 14^; large-scale transcriptomics studies for many *Drosophila* cell lines, developmental stages, and tissues ^15–17^; large-scale studies of transcriptional regulation^18^; and single-cell RNAseq analysis^19, 20^.

Protein-based resources and datasets provide an important complement to other ‘omics’ resources but are hindered in scale by technological challenges. Nevertheless, several efforts have generated physical and data resources relevant to *Drosophila* proteins. The first attempt at generating a binary protein interactome map for *Drosophila* at proteome-scale was released two decades ago^2^, followed by a few attempts at smaller scale^21–23^. In addition, the large-scale *Drosophila* Protein Interaction Map (DPiM) project, which used affinity purification followed by mass spectrometry (AP-MS), identified associations for ∼5,000 fly bait proteins^24^, a project that was made possible by the systematic ORFeome cloning project of the Berkeley *Drosophila* Genome Project (BDGP)^25^. Moreover, databases of known and predicted *Drosophila* PPIs have been established and updated, such as the Drosophila Interaction Database (DroID)^26–28^ and databases with multi-species coverage, including the Molecular Interaction Search Tool (MIST)^29^, BioGRID^30^, and IntAct^31^. Nevertheless, discovery of high-confidence binary interactions using ORF collections and up-to-date methods has remained limited in *Drosophila*.

To address this unmet need, we applied to *Drosophila* the overall strategy for large-scale, high-confidence detection of binary protein interactions and data integration that was recently reported for the human proteome^4^. Our approach involved two distinct configurations of the Y2H assay for a total of four all-by-all Y2H screens of 10,000 x 10,000 *Drosophila* proteins and resulted in a new *Drosophila* binary interaction dataset, the “FlyBi” dataset, of 8,723 binary interactions among 2,939 proteins. Subsequent reanalysis of previous datasets and integration of FlyBi data and literature-based binary interactions of comparable quality resulted in an expanded, high-confidence *Drosophila* reference interaction (DroRI) network of 17,232 binary interactions among 6,511 proteins. We tested the utility of the data to predict function by generating a putative autophagy interaction network that we validated *in vivo* using autophagy-related assays. The ORF clone collection and data resources generated in this project, are available from multiple public sources, provide a foundation for additional proteomics studies and for the generation of new hypotheses regarding protein functions in *Drosophila*.

### Expansion of the binary protein-protein interaction network for *Drosophila*

Performing Y2H screens with multiple, state-of-the-art versions of the assay can lead to increased high-quality coverage of binary interactions, as demonstrated by analyses that use existing knowledge as a benchmark for quality analysis (e.g., see^4^). We have demonstrated that both specificity and sensitivity of maps can be increased by improving any one of the four parameters of the ‘empirical framework’: i) completeness of the search space to be explored; ii) assay sensitivity; iii) sampling sensitivity; and iv) precision^32–35^. Since the *Drosophila* proteome contains ∼13,900 confirmed or predicted protein-coding genes, the complete search space to be eventually explored is at least a 13,900 x 13,900 matrix of 1.9×10^8^ combinations. The first systematic attempt was performed by screening ∼10,000 baits against two cDNA libraries and a pool of ∼10,000 ORF clones^2^. A significant limitation of that study was that at the time it was not feasible to sequence the full collection, such that the identities of all bait-prey pairs tested are not known. In addition, which partial or full-length isoforms were tested is not known; only a single assay version was used; a limited number of replicate screens were performed; gene annotations were of poorer quality than they are now; and the precision of the assays was heterogenous, with a subset of 4,780 PPIs reported as reaching acceptable quality levels^2^.

To improve our knowledge of the *Drosophila* binary interactome network, we chose to perform four large-scale Y2H screens with a set of ∼10,000 *Drosophila melanogaster* proteins of known sequence (see https://www.fruitfly.org/DGC/index.html and ^36^): two screens in each of two different configurations differing in the position of the Gal4 activation domain (AD) fusion, *i.e.* N- or C-terminus, and in the overall level of exogenous expression, *i.e.* using either centromeric or two-micron based expression vectors^4^. These four “all-by-all” screens represented 400 million combinations of protein pairs (**Fig. 1A** and **Suppl. Fig. 1**). First pass pairs (FiPPs) identified in the primary Y2H screens were systematically retested in pairwise tests, followed by sequence confirmation. The resulting list of sequence-confirmed putative binary interactors was supplemented by using a computational network-based approach to predict additional interaction pairs based on pairs in the assay version 1 screens followed by experimental validation (see **Fig. 1A**, Methods, and ^37^), altogether resulting in 332 experimentally predictions confirmed in the pairwise test.

**Figure 1:**
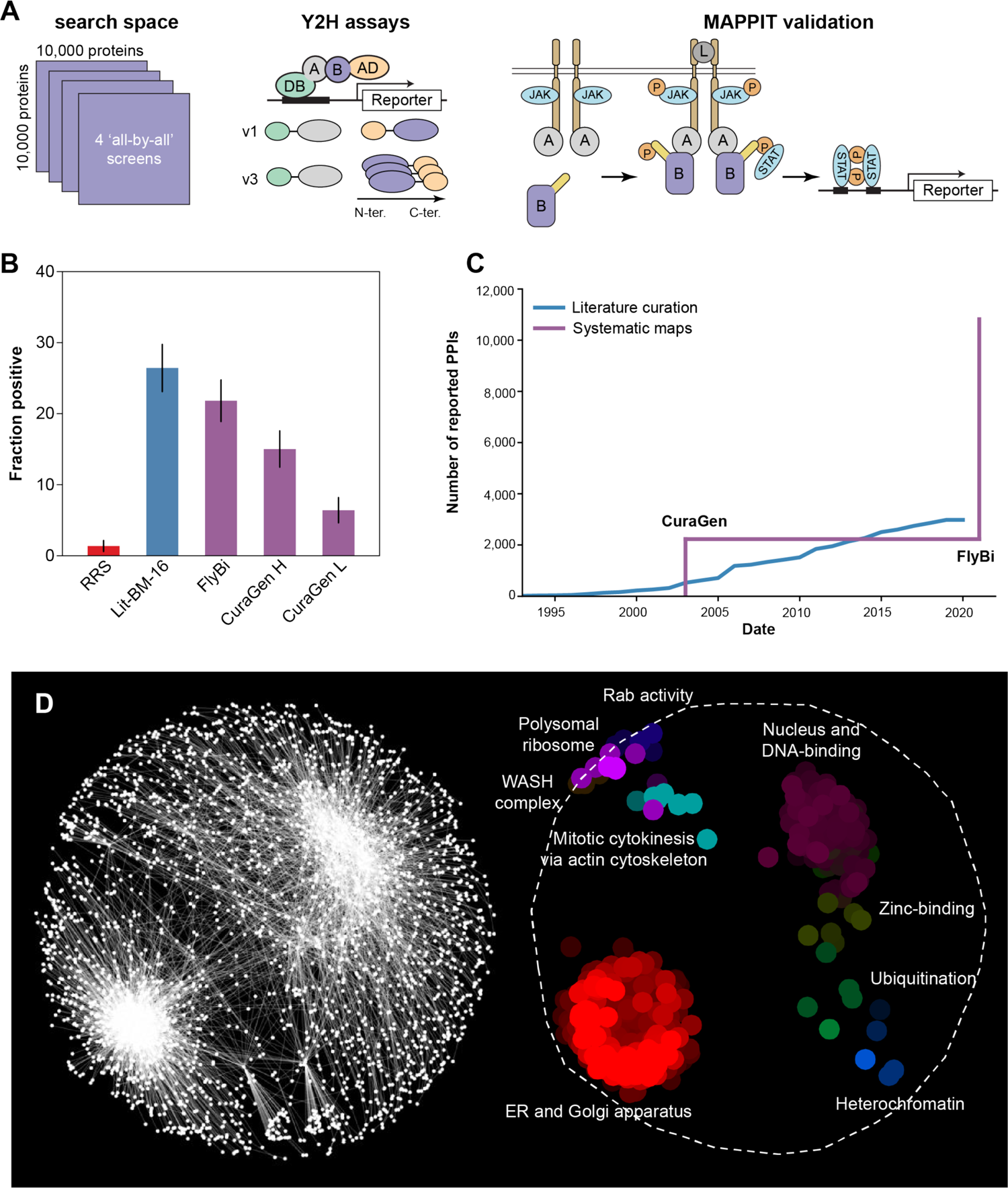
Large-scale, all-by-all binary interaction screens of 10,000 *Drosophila* proteins. **A.** Schematic of the systematic screening pipeline. Left, the search space covered and the Y2H assay versions used. Center, assay versions used in the Y2H screen. Right, MAPPIT validation assay. **B.** Fraction of pairs positive in the MAPPIT validation assay for the following sets: random reference set (RRS; red), literature-curated binary pairs with multiple evidence (at the time of the assay; Lit-BM-16; blue), FlyBi pairs, and CuraGen pairs at high (H) and low (L) cutoff values as defined by Giot, Bader et al. 2003 (purple) **C.** Total number of binary interactions in literature and systematic interactome maps over the past 20 years. Blue line, the total number of binary interactions accumulated in the literature (Lit-BM), as curated in FlyBase and displayed based on date of publication. Purple line, the number of interactions from systematic interactome mapping effort based on the date of public release of systematic binary datasets. **D.** Network-based spatial enrichment analysis (SAFE) results for the FlyBi dataset. Clusters of genes enriched for gene ontology (GO) terms are highlighted.

As an initial test of the quality of the experimental and computational binary protein pairs, we established (i) a small, high-confidence positive reference set (PRS), (ii) a random reference set (RRS) of the same size (**Suppl. Table 1**), (iii) a larger list of literature-curated binary pairs for which multiple lines of evidence support the interaction (at the time of the analysis, i.e., Lit-BM-16, **Suppl. Table 2**, or the most recent available, i.e., Lit-BM-20, **Suppl. Table 3**), and (iv) a list of literature-curated binary pairs for which only one line of evidence supports the interaction (Lit-BS, **Suppl. 4**), similar to what was done for the human reference interaction (HuRI) network^4,38^. The pairwise testing rates from the computational predictions based on the first and second screen as an input (L3; see Methods) were 90±10% for the top 100 predictions, 80±5% for the top 500 predictions, and 71±3% for the top 1000 predictions. These remarkable precision values indicate strong, highly non-random network patterns even in just one screen and open up the possibility of extending the experimentally obtained high-throughput interactome using computational predictions followed by pairwise tests. Pairs curated from the literature were recovered at lower rates (13±3% for Lit-BM-20 pairs).

Altogether, we identified 1,726 interaction pairs in assay version 1, screen 1; 1,029 in assay version 1, screen 2; 3908 in assay version 3, screen 1; and 3509 in assay version 3, screen 2; and we supplemented these with the 332 confirmed pairs based on computationally prediction, resulting in a total FlyBi dataset of 8,723 unique interaction pairs for 2,939 genes. The FlyBi dataset of interaction pairs is available as **Suppl. Table 5** and the FlyBi project webpage (http://flybi.hms.harvard.edu/). In addition, these pairs have been integrated with other datasets at IntAct (https://www.ebi.ac.uk/intact/)^39^ and in MIST (https://fgrtools.hms.harvard.edu/MIST/)^29^.

### Quality analysis using the MAmmalian Protein-Protein Interaction Trap (MAPPIT) assay

Verifying putative binary interactions with orthogonal assays provides a method for quality analysis that can be used to define cut-off values prior to integration of data from different sources. Thus, our next goal was to analyze the quality of the FlyBi pairs and of Lit-BM pairs (available at the time of the experiments; Lit-BM-16; see **Suppl. Table 2**), as well as binary pairs from the literature with only a single piece of evidence (Lit-BS), pairs identified in the previous large-scale *Drosophila* Y2H study (CuraGen pairs)^2^, binary pairs identified in additional Y2H studies made available at DroID^27, 28^, and interactions identified in the DPiM project^24^. We experimentally tested randomly selected subsets of pairs from the FlyBi and other datasets using the MAmmalian Protein-Protein Interaction Trap (MAPPIT) assay^40^. With the MAPPIT assay, binary interactions between two proteins expressed in mammalian cells activate signaling by an otherwise inactive cytokine receptor. The lists of 1,941 FlyBi, 209 Lit-BM-16, and 188 Lit-BS pairs tested in the MAPPIT assay are provided as **Suppl. Table 6** and **Suppl. Table 7**, and the subset of 323 pairs that were positive in the assay is provided as **Suppl. Table 8**.

The results of this analysis made it possible for us to apply a cut-off value for CuraGen pairs that produced a list of pairs of equivalent high quality as compared with FlyBi pairs from assay version 1 (N-N terminal configuration) as judged by performance in this orthogonal MAPPIT assay. The Giot et al. study reports that 4,780 interactions among 4,679 proteins met the cut-off value of 0.5 for high-confidence as defined in that study^2^. We found that CuraGen pairs with a confidence score of 0.7 or higher as defined in ^2^ have a similar recovery rate in the MAPPIT as compared with FlyBi pairs (**Suppl. Figs. 2 and 3**). Thus, a total of 2,232 protein pairs from the CuraGen dataset met the quality cut-off criteria for integration into our final reference map as described below. We note that pairs from the FlyBi assay ‘version 3’ screens (N-C terminal configuration) did not validate as well as the ‘version 1’ screen pairs (N-N terminal configuration) (**Suppl. Fig. 3**). Literature pairs also did not validate at the same rate when tested using the C-terminal version of MAPPIT. We attribute this to the fact that the MAPPIT assay has not been optimized for screens performed using C-terminally-fused ORFs. As expected due to significant differences in the assay formats, assay types, and other relevant factors, the positive rates with MAPPIT were lower for DPiM^41^ and for the group of previous smaller-scale Y2H studies available from DroID^27, 28^ (**Suppl. Fig. 3**). These other studies contributed to defining the Lit-BM, e.g., as the source of additional evidence for some pairs, and notably, the Lit-BM performs significantly better than the Lit-BS (**Suppl. Fig. 3**). This provides one indicator among many that these other datasets have clear value as part of an effort to fully document PPIs in *Drosophila*.

**Figure 2:**
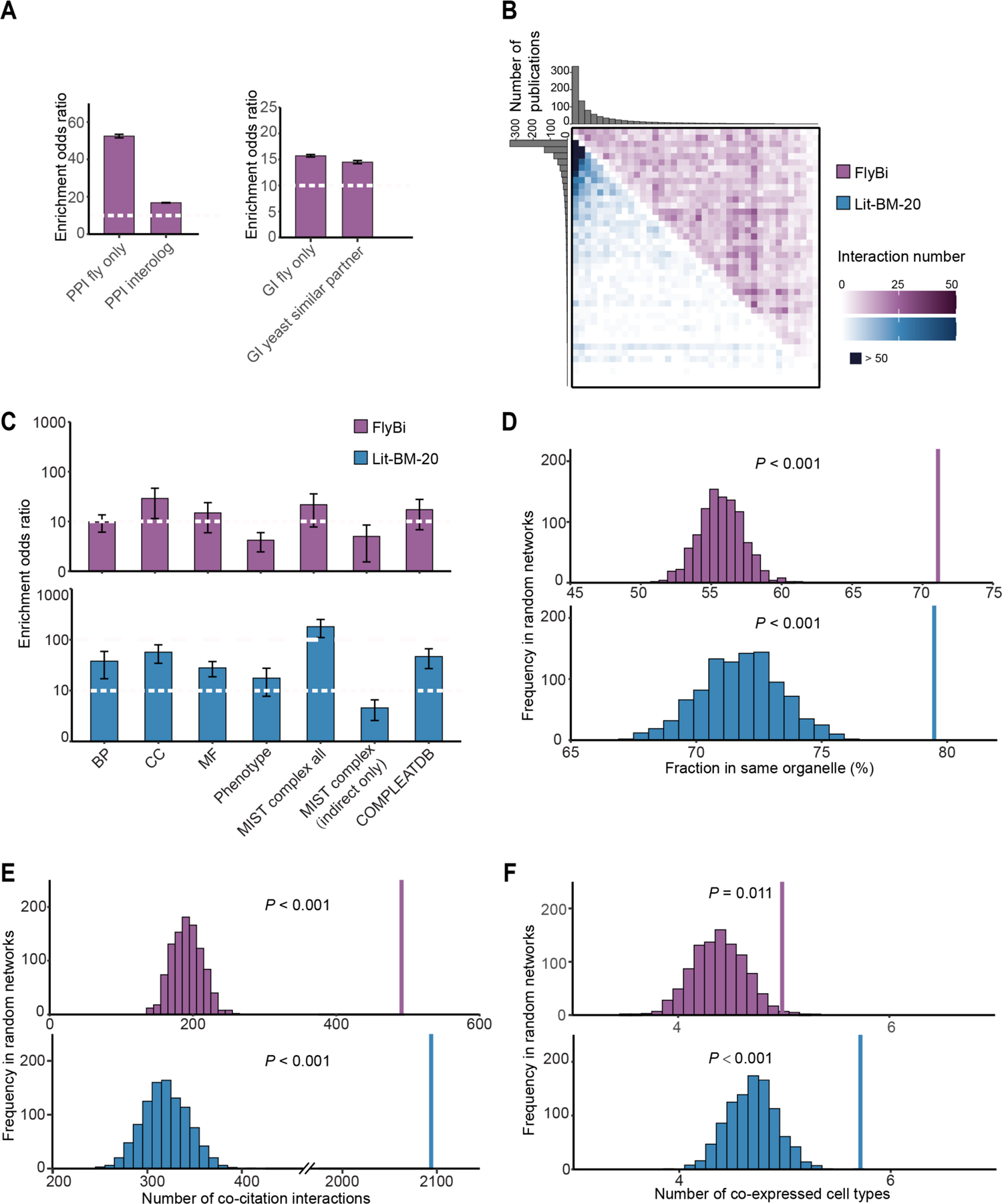
Bioinformatics analysis of the FlyBi Y2H dataset. **A.** Comparison on FlyBi with protein-protein and genetic interactions (PPIs and GIs) as annotated in the MIST database. The fraction of FlyBi pairs that overlap with published *Drosophila* PPIs or interologs (i.e. putative PPIs mapped based on orthologous genes in other major model organisms) were analyzed by comparison to 1,000 random networks generated by shuffling the nodes. The FlyBi dataset shows significant enrichment for published PPIs. Overlap with *Drosophila* GIs and with gene pairs with similar genetic interactors in yeast were also analyzed. **B.** Adjacency matrix for binary interactions reported in the literature with multiple lines of evidence (Lit-BM) and FlyBi interactions blue and red, respectively. For the visualization, proteins were binned and ordered along both axes based on the number of corresponding publications. The color intensity of each square reflects the total number of interactions between proteins in the corresponding bins. C-F: Biological significance analyses. Red, FlyBi; blue, Lit-BM. **C.** Enrichment of binary interactome maps for functional relationships and co-complex memberships. BP, biological process; MF, molecular function; CC, cellular component; Phenotype, shared phenotypes in *Drosophila* as annotated by FlyBase; MIST complex all, all annotated indirect interactions in MIST (might or might not be supported by direct evidence); MIST complex only indirect, all interactions annotated as supported only by indirect evidence in MIST; COMPLEATDB, complex annotations (only literature-based complexes were selected). The dashed white line provides a visual reference point for comparison of bar heights. For **D,E** and **F**, the single bar on the right shows the result for FlyBi data (purple) or Lit-BM (blue), and the multiple bars to the left show results for 1,000 randomized networks. **D.** Co-localization analysis. Shown, the fraction of the interacting partners that share the same organelle annotation, as compared with results for 1,000 randomized networks. Organelle annotations as predicted by PSORT and DeepLoc. **E.** Co-citation analysis. Shown are the numbers of interacting partners that are cited in the same publication(s), as compared with results for 1,000 randomized networks. Only publications associated with fewer than 100 genes were considered. **F**. Co-expression analysis. Shown is the average number of co-expressed cell types defined by cluster analysis of a single-cell RNAseq dataset for the *Drosophila* midgut, as compared to results with 1,000 randomized networks.

**Figure 3:**
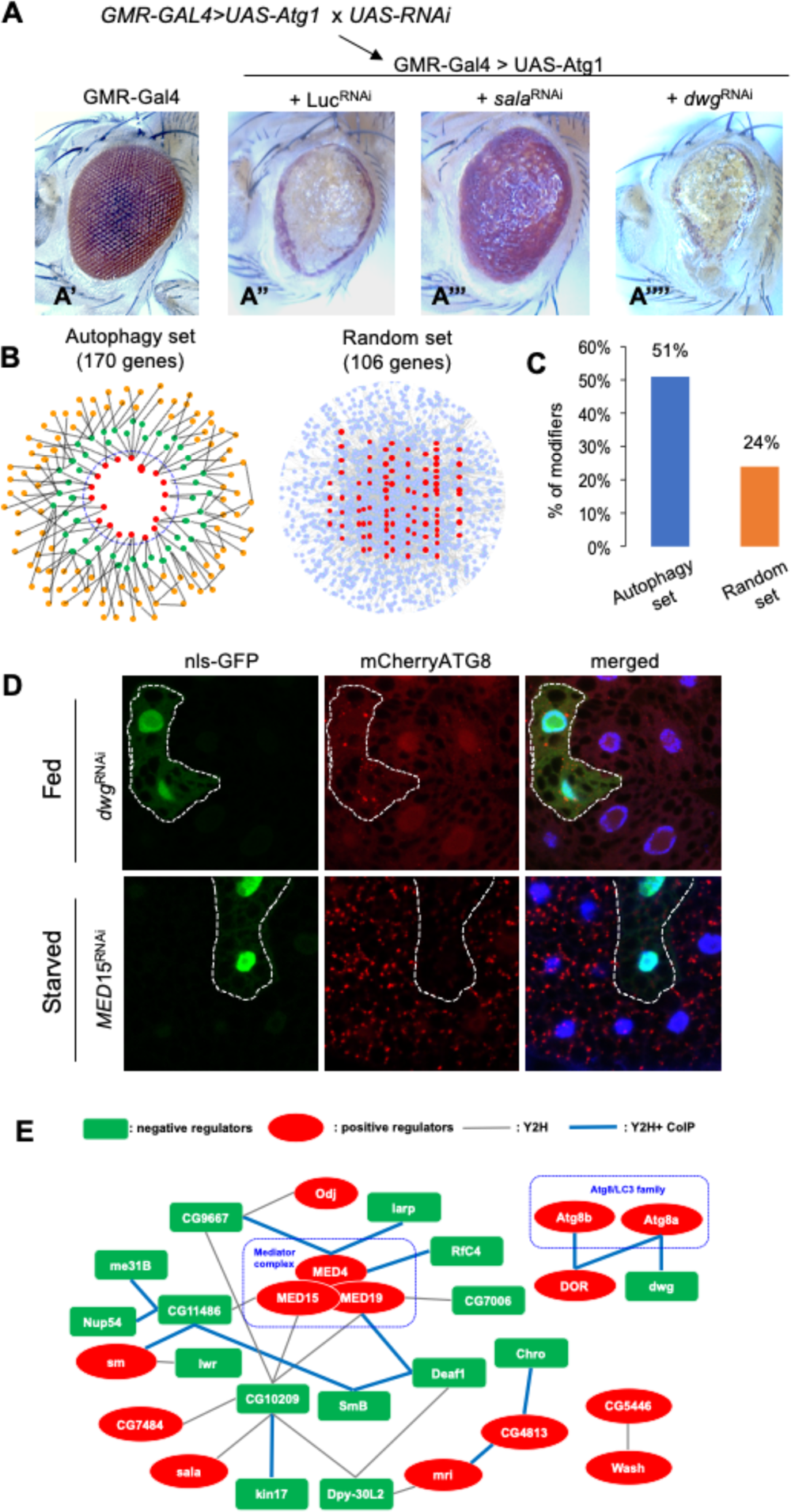
Identification of an autophagy regulatory network using the FlyBi dataset. **A.** Genetic cross and example phenotypes for RNAi knockdown in the presence of *Atg1* over-expression. Two sets were compared: an experimental set defined based on predicted interaction in the FlyBi dataset with known autophagy components or their interactions (**Suppl. Table 9**) and a randomly selected set. **A’-A’’’’.** Representative adult *Drosophila* eye phenotypes from control and experimental assays for modification of the *Atg1* overexpression phenotype. **A’**, Gal4-only control. **A’’**, Ectopic expression of *Atg1* using the eye-specific *GMR-GAL4* driver results in a rough eye and reduced eye size. The effect is reduced in the presence of *Sala^RNA^*^i^ (**A’’’**) and more severe in the presence of *dwg^RNAi^* (**A’’’’**). **B.** Visualization of the *in vivo* functional validation of autophagy-related interactors described in A. Pink indicates putative enhancers. Green, putative suppressors. Yellow, inconsistent results with multiple RNAi lines. Grey, no effect on the *Atg1* over-expression phenotype. Red edge, overlap with published datasets. Grey edge, novel interactions. **C.** Percentage of RNAi lines that behaved as putative genetic modifiers of *Atg1* over-expression. **D.** Distribution of mCherry-ATG8a in the larval fat body (fed or starved conditions). Clonal expression of *dwg^RNAi^* in GFP-labeled cells induced the formation of mCherry-ATG8a puncta under fed condition while *MED15^RNAi^* abrogated starvation-induced Atg8a puncta. **E**. Putative autophagy regulator network based on knockdown, FlyBi data, and co-immunoprecipitation (co-IP) data (see **Suppl. Fig. 5** and **Suppl. Fig. 6**). Green, putative suppressors of autophagy; red, putative inducers of autophagy. Grey edges, direct interactions as reported in the FlyBi dataset. Blue edges, interactions reported in FlyBi and confirmed by co-IP. Of the genes In the network, 6 (30% of total) computed relatively unstudied genes (‘CGs’) were added to the network by our studies.

### Comparison of FlyBi interactions with existing knowledge

We next compared FlyBi pairs with interaction data from a variety of data repositories that are integrated in MIST^29^ (**Fig. 2**). We generated 1,000 randomized versions of the FlyBi network by node shuffling. Interacting pairs in the FlyBi dataset show significant overlap with physical interaction data obtained from previous studies in *Drosophila* and physical interactions mapped from orthologous genes (‘interologs’) (**Fig. 2A**). We also observed some overlap between FlyBi binary interaction pairs and genetic interaction (GI) data for *Drosophila* and between orthologs of fly genes in the budding yeast *Saccharomyces cerevisiae* (**Fig. 2A**). To further analyze FlyBi interactions, we determined the count of literature citations for each gene in the Lit-BM-20 or FlyBi dataset. As expected, interactors in Lit-BM-20 are biased towards well-studied genes (i.e., genes with larger numbers of literature citations). By contrast, we did not observe this bias for genes in the FlyBi dataset (**Fig. 2B**), consistent with the large-scale, all-by-all approach we took to generate the data. We next compared gene ontology (GO) annotations for the two proteins in each pair in three categories—biological process, molecular function, and cellular component—as well as phenotype annotations from FlyBase. For both Lit-BM-20 and FlyBi pairs, we observe significant enrichment for genes with the same GO and/or phenotype annotations as their interacting partners. Moreover, the level of enrichment as compared with random controls is comparable for Lit-BM-20 and FlyBi pairs (**Fig. 2C**). We also compared binary interactions with protein complex-based interaction data, and with components of protein complexes as annotated in literature^42^, and observed enrichment in both the Lit-BM-20 and FlyBi sets (**Fig. 2C**). In addition, interacting proteins identified in our study are more likely to be found in the same organelle and in the same cell type, as well as reported in the same publications, compared to the random controls (**Fig. 2D,E,F**).

Comparing the Lit-BM-20 and FlyBi sets to random networks reveals that FlyBi interaction pairs are of a quality that is comparable to the high-confidence published binary interactions that make up the Lit-BM-20 and are less biased. As such, these sets can appropriately be combined to generate a high-quality *Drosophila* reference interaction (DroRI) network. We built a new, high-confidence DroRI network by integrating the FlyBi data, CuraGen data that meet the cut-off for data quality equivalent to the FlyBi data, and all other high-confidence binary interactions (i.e., literature-based interactions). The DroRI network is comprised of 17,232 interactions among the protein products of 6,511 genes and can be queried and accessed at a dedicated page at MIST. To facilitate integrated data mining and hypothesis generation, we integrated tissue-specific bulk RNA-seq data generated by the modENCODE consortium^43^ into MIST. This makes it possible for users of MIST to project any of the tissue-specific transcriptomics datasets onto the DroRI network, and reveal the subset of network interactions predicted to occur in a given tissue (**Suppl. Fig. 4**).

We also compared the DroRI network with interaction data from the mass spectrometry-based DPiM study, and with binary interaction maps from other species. Given the differences in coverage and approach, we expected the overlap between DroRI and these other datasets to be modest. Consistent with that expectation, we found that among the 17,232 binary interactions in the DroRI network, only 54 pairs overlap with DPiM data and 2,661 pairs overlap with the complete set of interactions detected by mass spectrometry as annotated in MIST. With regards to other species, we find that comparison of the DroRI and HuRI networks reveals 714 of a total of 9,332 interactions for which both orthologs are present in both datasets are identified as binary interactors in both networks. The total set of DroRI binary interactions for which orthologs are detected as binary interactors in any of the species included in MIST (human, rat, mouse, zebrafish, *X. laevis, X. tropicalis, C. elegans, S. cerevisiae,* and *S. pombe*) is 1,355. The low level of overlap likely also reflects differences in what is discoverable using different types of assays, whether or not the correct isoform is being tested, other sources of false negative discovery, and meaningful biological differences.

Ultimately, the value to the *Drosophila* research community of the DroRI network, and the new FlyBi dataset in particular, will be revealed by exploring its use, such as for the development of new hypotheses regarding protein function. We describe the results of one such exploration below.

### Generation of an autophagy network using FlyBi data

To experimentally test the predictive value of interactions represented in the DroRI network and in particular, to test the predictive value of the new FlyBi binary interaction pairs with regards to shared gene function, we chose to focus on autophagy. Autophagy has been extensively studied in multiple species^44^; has been characterized using protein-centric approaches in human cells^45^; is a conserved process with human health relevance^46^; and is easily studied *in vivo* in *Drosophila* using multiple well-established assays^47, 48^. To identify new regulators of autophagy, we used FlyBi data to define a list of candidate autophagy-related proteins and a control set. To build a putative autophagy-related list, we first assembled a list of 19 known autophagy regulators (List 1 in **Suppl. Table 9**). Next, we mined the FlyBi data for interactors with these autophagy regulators and identified 48 candidate interactors (List 2 in **Suppl. Table 9**). One of the 19 known regulators we included is Atg8a. There are five interactors with Atg8a in the FlyBi dataset. We note that two of these five were also identified in a recent Y2H screen for interactors with Atg8a^49^; i.e., Diabetes and obesity regulated (DOR), a known autophagy regulator, and CG12576, an uncharacterized protein. To expand the candidate list, we again mined the FlyBi data, and identified 103 additional potential interactors of List 2 proteins (List 3 in **Suppl. Table 9**). By combining Lists 1, 2, and 3, we generated a putative PPI network related to autophagy that includes four core autophagy-related genes and 166 candidates (170 gene ‘autophagy set’) (**Suppl. Table 10**). As a control set, we chose at random 106 genes from the FlyBi dataset (‘random set’) (**Suppl. Table 10**).

To test for autophagy-related functions, we performed loss-of-function experiments using RNAi (**Fig 3A**) combined with overexpression of *Atg1*, which encodes a protein kinase essential for autophagy (**Fig. 3A**). Overexpression of *Atg1* in the *Drosophila* eye induces a high level of autophagy, leading to a rough eye phenotype (**Fig. 3A**, compare A’ and A’’)^50, 51^. To test the roles of the ‘autophagy set’ genes, we determined if RNAi knockdown of these candidates modified the *Atg1*-induced rough eye phenotype. A total of 477 RNAi lines targeting 166 genes were tested in a *GMR-Gal4>UAS-Atg1* background. We found that 234 lines, corresponding to 137 genes, modified the severity of the *GMR-Gal4>UAS-Atg1* phenotype (**Fig. 3B** and **Suppl. Table 10**). To address whether the data from FlyBi used to generate the autophagy network helped enrich for potential autophagy components, we randomly selected 106 genes from FlyBi as a control list of comparable size (‘random set’) (**Fig. 3B**) and tested these genes in the *GMR-Gal4>UAS-Atg1* assay. Altogether, 26 of the lines tested (24%) modified the severity of *GMR-Gal4>UAS-Atg1* phenotype (**Suppl. Table 10**). We tested multiple lines per gene in the autophagy set and only a single line per gene in the control set. Thus, to appropriately compare the percentage of modifiers between the autophagy set and the control set, we randomly selected one RNAi stock per gene from the autophagy set five times, generating five independent data sub-sets (see Methods) (**Suppl. Table 10**). RNAi lines tested in the autophagy sub-set modified the *GMR-Gal4>UAS-Atg1* phenotype in 50%, 52%, 47%, 51%, 55% of cases (average = 51%), compared to 24% in the random set (**Fig. 3C**). Altogether, these results indicate that the targeted candidate gene screen approach is more efficient at identifying new potential modifiers of autophagy-related functions.

We next tested putative autophagy regulators identified in the *GMR-Atg1* screen using a different assay performed at a different lifecycle stage and in a different tissue (**Fig. 3D**). This assay interrogated autophagy-related processes in the larval fat body, a nutrient storage organ analogous to the human liver in which autophagy is quickly induced by starvation^52^. Under fed conditions, expression of the autophagosomal marker mCherry-ATG8a shows diffuse localization throughout the cells. Upon starvation, mCherry-ATG8a redistributes to form punctate structures (autophagosomes) in the cytoplasm. Of the 234 RNAi lines identified in the *GMR-Gal4>UAS-Atg1* eye screen, 60 (26%) increased fat body Atg8a puncta under fed conditions, while 41 lines (18%) inhibited fat body Atg8a puncta formation upon starvation (**Suppl. Table 10**). As an example of a negative autophagy regulator, depletion of *dwg* in GFP-labeled flip-out clones induced Atg8a punctate formation under nutrient rich conditions (**Fig. 3D**, top panels). In addition, as an example of a positive regulator of autophagy, depletion of *MED15* suppressed starvation-induced Atg8a puncta formation (**Fig. 3D**, bottom panels). Altogether, our candidate gene approach allowed us to quickly and efficiently identify new genes that are likely to be regulators of autophagy.

In total, 101 RNAi lines corresponding to 66 genes from the ‘autophagy set’ list were able to modify Atg1-induced eye defects and alter Atg8 puncta. Of the 66 genes, we chose 39 genes for which there are at least two RNAi lines available and results with both lines had consistent effects in both the fat body and the eye phenotype screens. We tested whether components of the autophagy network can physically interact in *Drosophila* cells by co-immunoprecipitation (Co-IP). We expressed Flag- and GFP-tagged proteins together in *Drosophila* S2R+ cells, pulled down GFP-tagged proteins, and determined whether they were associated with Flag-tagged proteins (**Suppl. Fig. 5, Suppl. Fig. 6, Suppl. Table 11**). Of the 39 genes, one of the GFP-tagged proteins, CG10209, was expressed at very low levels. To overcome this issue, we designed a smaller Flag-tagged form of CG10209 and enriched it using Flag-beads (**Suppl. Fig. 6**). Of 29 pairs we tested, an interaction was detected using co-IP for 16 (55%), providing support for the high quality of the FlyBi dataset **Fig. 3E**.

### Dwg suppresses autophagy by binding to insulator elements near *ATG* genes

One of our candidates, *deformed wings (dwg;* also known as *Zw5),* encodes a *Drosophila* insulator protein responsible for enhancer blocking and support of distant interactions, contributing to the organization of chromosome architecture^53^. Our genetic test showed that *dwg* is a putative negative regulator of autophagy. Consistent with this, whole larval lysate of *dwg* mutants showed a higher level of autophagy, indicated by increased lipidated Atg8a (Atg8a-II) compared to lysate from control (**Fig. 4A**).

**Figure 4:**
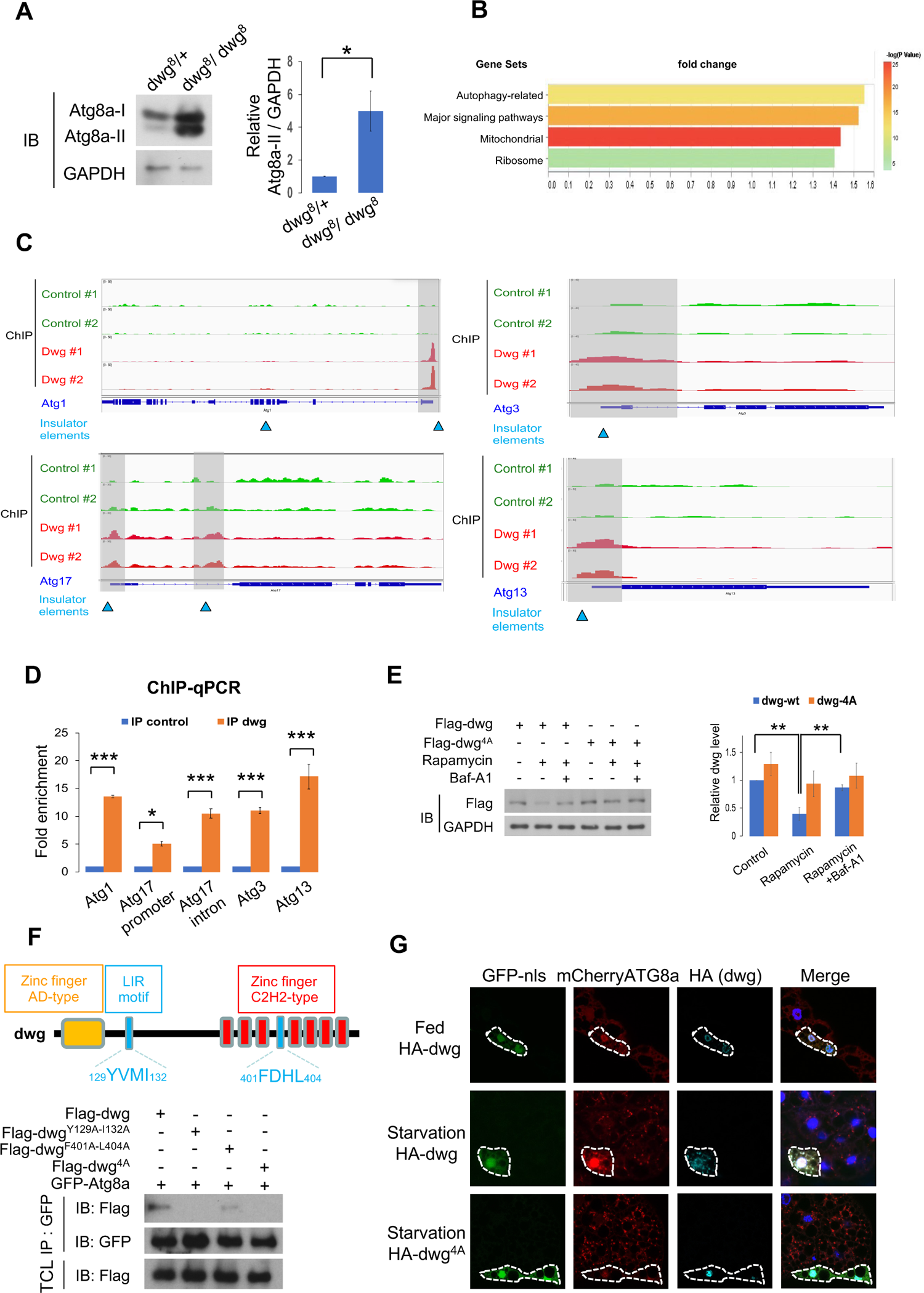
Dwg is both a negative regulator and a substrate of autophagy. **A.** Flies homozygous mutant for a loss of function allele of *dwg* exhibit increased levels of autophagy. Control (*dwg*^8^/+) and *dwg* mutant (*dwg*^8^/*dwg*^8^) larva were subjected to immunoblotting with antibodies as indicated. Measurements shown are mean ± SEM of triplicate experiments. Significance was determined by t-test. *p<0.05. **B.** Gene group enrichment analysis of analyzed ChIPseq data. Bar length, fold change in enrichment. Colors, strength of significance (p-value of -log10 for each term). C-D, Dwg binds to insulator regions of *Atg* genes. Example browser images for *Atg1, Atg3, Atg13,* and *Atg17* from ChIP-seq experiments in S2R+ cells expressing Flag-Dwg. Aggregate data from two independent ChIP-seq experiments are shown. Insulator binding regions are indicated with blue triangles. **C.** Dwg occupancy at or near insulator regions of *Atg* genes as revealed by ChIP-qPCR. **D.** One-Way ANOVA followed by Tukey’s multiple comparison test was performed to identify significant differences; data shown as means ± SEM of three independent experiments; ***P<0.001, *P<0.05. **E.** Autophagic activity regulates Dwg protein levels. S2R+ cells transfected with *Flag-dwg or - dwg^4A(Y129A-I132A-F401A-L404^*)) were treated with Rapamycin, an autophagy inducer, or Bafilomycin A1 (Baf-A1), a lysosomal inhibitor. Dwg and GAPDH protein levels were analyzed by immunoblotting (IB) with antibodies as indicated and quantified. One-Way ANOVA followed by Tukey’s multiple comparison test was performed to identify significant differences; data shown as means ± SEM of three independent experiments; **P<0.01. **F.** Mapping Dwg-Atg8a interaction sites. Schematic presentation of domain structures and LIR (LC3-interacting region) motifs of Dwg. S2R+ cells transfected with *GFP-Atg8a, Flag-tagged dwg,* or *Flag-tagged dwg* with different mutations in LIR motif (*dwg^Y129A-I132A^*, *dwg^F401A-L404A^,* or *dwg^4A(Y129A-I132A-F401A-L404^*)) for 48 hr followed by immunoprecipitations with anti-GFP nanobody. The immunoprecipitated proteins and total cell lysates were analyzed by immunoblotting with antibodies as indicated. **G.** Disruption of the Dwg-Atg8a interaction inhibits autophagy. Clonally expressed HA-tagged Dwg is located in the nucleus under fed conditions. Upon starvation, it is detected in the cytoplasm and co-localizes with autophagosomes as labeled by mCherry-Atg8a. Versions of Dwg with mutations in Atg8a-binding sites (*dwg*^4A^), however, remains in the nucleus and inhibits autophagy. Fat body cells were stained with DAPI.

We hypothesized that as an insulator, the Dwg protein might regulate autophagy through binding to insulator elements on chromatin and blocking enhancer functions. We therefore performed chromatin-immunoprecipitation followed by next-generation sequencing (ChIP-seq) to identify Dwg downstream targets. Gene group enrichment analysis revealed that the chromatin regions of autophagy-related genes and genes related to mitochondria, major signaling pathways, and ribosomes are targeted by Dwg (**Fig. 4B**). Interestingly, the Dwg-binding regions verified by ChIP-qPCR are located at or near insulator elements in four core ATG genes, Atg1, Atg3, Atg13, and Atg17 (**Fig. 4C,D**)^54^. Dwg can suppress enhancer functions, thus leading to inhibition of transcription^55^. Consistent with this, we also observed that *dwg* mutants showed higher mRNA expression of *ATG* genes (**Suppl. Fig. 7**). Taken together, these results suggest that Dwg binds to the insulator elements present in the *ATG* genes, presumably suppressing their transcription.

### Dwg is subjected to autophagy-lysosomal degradation

Autophagy is considered a highly selective pathway that targets specific substrates for degradation and selectivity is thought to rely mainly on the interaction between LC3/ATG8 family proteins and cargo/adaptor proteins^56^. Interestingly, our co-IP results suggest that Dwg physically interacts with Atg8a (**Suppl. Fig. 8**) and FlyBi data further suggest that the interaction is direct. These results suggest that Dwg is a substrate for autophagy. To test this hypothesis, we expressed Dwg in S2R+ cells and treated cells with an autophagy inducer (Rapamycin) or a lysosomal inhibitor (Bafilomycin A1; BafA1). Immunoblots revealed that Dwg protein levels were reduced following Rapamycin treatment, whereas the Rapamycin-induced reduction of Dwg protein can be reversed by cotreatment with Bafilomycin A1 (**Fig. 4E**), indicating that Dwg is degraded by autophagy.

The mammalian ortholog of Atg8a, LC3, interacts with LIR (LC3-interacting region) motifs, W/F/Y-x-x-L/I/V, on substrates for autophagic degradation^56^. There are four potential LIR motifs predicted in Dwg (**Suppl. Fig. 8**)^57^. To characterize which LIR motifs are responsible for interactions with Atg8a, we generated four Dwg deletion mutants, Dwg-F1-F4. Each one of them contains an individual LIR motif (**Suppl. Fig. 8**). Our co-IP results showed that Atg8a interacts with Dwg-F1 and Dwg-F4 (**Suppl. Fig. 8**), suggesting that it is the first and fourth LIR motifs that bind to Atg8a. Consistent with this result, Dwg with mutant LIR motifs (Dwg^Y129A-I132A^, Dwg ^F401A-L404A^, and Dwg^Y129A-I132A-F401A-L404A^ ^(4A)^) had dramatically reduced interactions with Atg8a, demonstrating that these two LIR motifs are Atg8a binding sites (**Fig. 4F**).

### Atg8a delivers Dwg from nucleus to autophagosomes for degradation

To elucidate the physiological role of the Dwg-Atg8 interaction, we expressed wild-type Dwg or Dwg with mutations in the two LIR motifs (Dwg^4A^) and examined their localization and effects in S2R+ cells and larval fat body. As expected, Dwg is localized in the nucleus (**Fig. 4G** and **Suppl. Fig. 9**). Inhibition of autophagosome degradation by Baf-A1 resulted in an increase of detectable Dwg in the cytoplasm and co-localization of Dwg with Atg8a punctae in S2R+ cells (**Suppl. Fig. 9**). Similarly, in the fat body, starvation induces translocation of Dwg to the cytoplasm, where it significantly colocalizes with autophagosomes (**Fig. 4G**). These results further support that Dwg is a substrate of autophagy. Importantly, expression of Dwg with LIR motif mutations was restricted to the nucleus and strongly inhibited autophagy in both S2R+ cells and fat bodies, suggesting that Atg8a is able to interact with and deliver Dwg to autophagosome for degradation (**Fig. 4 G** and **Suppl. Fig. 9**). Altogether, our results suggest that disruption of the Dwg-Atg8a interaction not only stabilizes Dwg protein, but also allows Dwg to bind to insulator elements which suppress transcription of *Atg* genes, leading to autophagy inhibition (**Suppl. Fig. 10**).

## Discussion

In this work, we applied state-of-the-art experimental approaches to binary interaction mapping, together with experimental and bioinformatics-based quality analyses, to generate a next-generation reference binary interactome for *Drosophila*. The outcomes of our large-scale efforts include (i) a collection of ∼10,000 *Drosophila* ORFs in a Gateway-system entry vector; (ii) a new high-confidence Y2H dataset, the FlyBi dataset, which is comprised of 8,723 binary interactions among 2,939 proteins; and (iii) an integrated *Drosophila* reference interactome, DroRI, which is comprised of 8,723 binary interactions among 2,939 proteins. Features that distinguish the FlyBi project from past efforts include the quality and coverage of the ORF collection on which we based our Y2H screens^25^; use of improved versions of the Y2H system^4^; prediction of new interactors using a computational approach^37^; use of an orthogonal approach to define cut-off values for confidence for FlyBi Y2H data, computational predictions, and previously reported data^40^; and integration and comparison of FlyBi data with existing PPI and other datasets to generate the DroRI resource that can be navigated using MIST^29^.

Several indicators point to the value and high quality of FlyBi interaction pairs. For example, proteins in FlyBi pairs are less biased towards well-understood proteins as judged by the number of publications per gene as compared to existing pairs (**Fig. 2B**), such that they provide an important supplement to existing interaction datasets. Moreover, FlyBi pairs also performed as expected for a dataset enriched in biologically meaningful associations (**Fig. 2C-F**). Nevertheless, the pairs defined in this work have little overlap with interactors as defined using mass spectrometry-based approaches (e.g., the DPiM dataset) or with binary interactors observed in other species, observations that likely reflect both experimental and biological differences. Ultimately, the value of identification of large-scale binary datasets for biological and biomedical research lies in the ability to use individual identified putative PPIs and/or integrated networks to build new hypotheses that lead to efficient detection of new functional findings with *in vivo* relevance. To test this, we explored potential new interactors of proteins previously identified as relevant to autophagy in *Drosophila*. We chose to focus on autophagy because this process is well-studied in multiple species and has human health relevance, and because well-established *in vivo Drosophila* assays related to autophagy were available.

Our approach was to start with known proteins of the autophagy pathway, identify potential PPIs based on the FlyBi data, and test these candidates for autophagy-related phenotypes in *Drosophila* using two different *in vivo* assays. We performed a focused RNAi screen for putative genetic modifiers of the mild phenotype associated with over-expression in the eye of *Autophagy-related 1 (Atg1)*^50, 51^. Using the positive hits from the *Atg1* modifier screen we determined the distribution of fluorescent protein-tagged Atg8a in the *Drosophila* fat body under fed and starved conditions^58^. Following this approach, we identified a high-confidence sub-network of putative autophagy regulators (**Fig. 3**) and found that Dwg both regulates and is regulated by autophagy, providing the first evidence of reciprocal regulation between autophagy and chromatin regulators. Importantly, these findings show that using our interaction network constructed with FlyBi data, allowed us to enrich for genes relevant to the process of autophagy. The ability to use binary interaction data to reduce the full set of *Drosophila* genes to a subset of high-confidence candidates prior to *in vivo* phenotypic analyses, which can be both time- and resource-intensive, will unquestionably accelerate future studies.

## Materials and Methods

### Generation of a large-scale ORF clone resource

The entry clone collection was generated from 11,687 BDGP cDNA gold clones ^36^ (see https://www.fruitfly.org/EST/gold_collection.shtml) using attB-tailed PCR. See below for detailed descriptions of primer design and PCR amplification. The PCR products were quality controlled by detection and sizing on agarose gels as follows. PCR products were loaded into 1% (w/v) agarose gels (3 g Agarose, 300 mL 1xTAE) and run in 1xTAE buffer with New England Biolabs 1 kb ladder. Gels were imaged using a BioRad GelDoc XR system. Band sizes were calculated using BioRad Quantity One software (version 4.6.9). High-quality PCR products were cloned into the pDONR223 expression and cloning vector using BP Clonase. See below for a detailed description of the cloning protocol. Clones were stored as glycerol stocks prior to their use to generate yeast expression clones.

#### Primer Design

PCR primers were designed using a custom Perl script and Primer3 release 0.9^59^. The parameters applied were as follows: PRIMER_OPT_SIZE, 22; PRIMER_MIN_SIZE, 18; PRIMER_MAX_SIZE, 30; PRIMER_OPT_TM, 60; PRIMER_MIN_TM, 40; PRIMER_MAX_TM, 95; PRIMER_OPT_GC_PERCENT, 50; PRIMER_MIN_GC, 0; PRIMER_MAX_GC, 100; PRIMER_EXPLAIN_FLAG, 1; PRIMER_MAX_POLY_X, 18; PRIMER_SELF_ANY, 30; PRIMER_SELF_END, 30. For most cases, the Perl script was invoked as follows: ./generateBOBSPrimers.pl -method open -primer_min_size 15 -primer_max_size 17 -vector pDONR223 - order <source_cdna_clone_id>. When that failed, the script was run as follows: generateBOBSPrimers.pl-method open -primer_min_size 15 -nomaxprimerlen -vector pDONR223 -order <source_cdna_clone_id>. For dicistronic cDNAs, the script was run as follows: ./generateBOBSPrimers.pl -method open -primer_min_size 15 (-primer_max_size 17 | -nomaxprimerlen) -vector pDONR223 -target <nucleotide sequence of CDS> -identification <gene name> -order <source_cdna_clone_id>. For dicistronic cDNAs, the CDS to which primers were designed was chosen manually and somewhat arbitrarily based on function and size, with preference given to the longer CDS.

#### PCR amplification of ORFs

Templates were inoculated from the Berkeley Drosophila Genome Project (BDGP) Gold Collection into 1.2 mL 2x YT medium with appropriate antibiotic (Chloramphenicol at 100 µg/ml final conc., or Carbenicillin at a final concentration of 100 µg/ml). Cultures were grown overnight (16-18 hr) at 37°C at 300 rpm. The overnight culture was diluted 1:10 with sterile water. PCR primers were purchased desalted and resuspended in Tris-EDTA (TE) buffer at 20 µM concentration from Invitrogen. Pairs of primers were combined and diluted with Milli-Q water to a concentration of 1.25 µM (each primer). PCR reactions were performed using 5 µL Phusion HF Buffer (5X concentration), 0.5 µL dNTP (10mM each, New England Biolabs), 5 µL primer pair mix (1.25 µM each primer), 3 µL template (1:10 cell dilution), 0.25 µL Phusion DNA Polymerase (New England Biolabs), and 11.25 µL sterile Milli-Q water, for a total reaction volume of 25 µL. Touchdown PCR ^60^ was performed with the following cycling parameters: 98°C for 1 minute; 5 cycles of (98°C for 10 seconds; 56°C to 46°C, decrease by 2°C each cycle; 72°C for 7.5 minutes); 15 cycles of (98°C for 10 seconds; 72°C for 7.5 minutes); 72°C for 10 minutes; 4°C hold.

#### BP Clonase reactions

BP Clonase reactions were performed in a total volume of 5µL, consisting of 1 µL 5X BP Reaction Buffer, 1 µl pDONR223 vector (75 ng/µL, uncut), 1 µL BP Clonase, and 2 µL of attB-tailed PCR product. BP reactions were incubated at 25°C for 18 hr. Immediately following incubation, 2 µL of the BP reaction was transformed into 10 µL of chemically competent *E. coli* DH5-alpha cells (prepared in-house). The mixture was incubated for 30 min on wet ice, heat shocked for 40 sec at 42°C, and incubated for 2 min on wet ice. Finally, 90 µl of SOC medium was added and the transformations were incubated for 1 hr, 225 RPM, 37°C in an orbital shaking incubator. The entire transformation reaction was inoculated into 1mL LB/spectinomycin (100 µg/mL) and incubated 16-18 hr at 37°C, 300 rpm. Reactions and transformations were performed in 96-well format in standard thermal cycler plates. Glycerol frozen stocks (15% glycerol) were made by mixing 50 µL glycerol (30%) with 50 µL overnight culture.

### Amplification of ORFs for transfer to expression vectors and sequence analysis

We used PCR to amplify the ORFs from the large-scale ORF collection to generate a product that was used for cloning into the yeast expression vectors (see below) and useful for sequence analysis. PCR was performed using individually indexed 96-well M13 forward primers (Life Technologies) and a non-indexed M13 reverse primer (5’-GTAACATCAGAGATTTTGAGACAC-3’). The same amount of each amplicon from each plate was pooled as a single sample. Samples from each entry plate were sequenced using the Illumina platform. Sequencing reads were deconvoluted to the individual well level based on a combination of the 96-well index and the Illumina sample index, and by alignment to ORF sequences. A clone was deemed ‘sequence confirmed’ if a majority of the reads from the well (> 10 reads) aligned to the expected ORF sequence. Only entry clones that were sequence confirmed were re-arrayed and used for further processing.

### Preparation of Y2H expression clones from the large-scale ORF clone resource

Using the M13 PCR product from the entry ORFs, we performed a LR reaction into pDEST-DB, pDEST-AD-*CYH2* (assay version 1) and pDEST-AD-AR68 (assay version 3) using Gateway Technology (Invitrogen). Attributes of these plasmids are summarized in **Suppl. Table 12**. The DNA was isolated using a liquid handling robot (Qiagen 96-well Miniprep). DB ORF fusions were transformed into yeast strain Y8930 and AD ORF fusions into yeast strain Y8800.

### Y2H auto-activator identification and removal

Prior to the screen, haploid DB ORFs were spotted on SC-Leu-His media to test for auto-activation of the GAL1::HIS3 reporter gene in the absence of an AD-ORF plasmid. DB ORFs that grew on SC-Leu-His were removed from the collection.

### Y2H Screening

Large-scale Y2H screens were performed using two assay formats^4^. For the first two screens (assay version 1), pools of 1,000 ORFs as preys in pDEST-AD-*CYH2* were screened against single pDEST-DB ORF baits. Both AD and DB are fused to the N-terminus of the ORF and expressed from yeast centromeric plasmids (**Fig. 1A**, center panel, “v1”). For screens 3 and 4 (assay version 3), we used preys in pDEST-AD-AR68, in which the AD is fused to the C-terminus of the ORF and plasmid copy number reflects use of a 2-micron origin instead of the yeast centromeric chromosome. We used assay version 1 prey constructs and tested these against same baits (**Fig. 1A**, center panel, “v3”). A detailed workflow is provided in **Suppl. Fig. 1B** and follows what was reported for^4^. Briefly, following inoculation of DB and AD ORFs in selective media and overnight culture, 10 ul of each DB was mated against 5 ul of a pool of 1,000 AD’s (kilopool). After an overnight incubation at 30°C in liquid rich medium (YEPD), 10 ul of the culture was transferred into synthetic complete media (SC) without leucine or tryptophan (SC-Leu-Trp) to select for diploids. The following day, the culture was spotted on SC-Leu-Trp-His+3AT solid media to select for diploids in which the *GAL1::HIS3* reporter gene was activated. In parallel, diploid yeast cells were transferred onto SC-Leu-His+3AT solid media supplemented with 1 mg/l cycloheximide (CHX) for assay version 1 or 10 mg/l CHX for assay version 3. After 72 hr incubation at 30°C and one additional day at room temperature, we picked colonies that grew well on 3AT plates and did not grow on CHX plates.

### Yeast colony sequencing

To identify both bait and prey proteins for thousands of positive colonies, we used a method called SWIM-seq (Shared-Well Interaction Mapping by sequencing) as described^4^. Briefly, DB and AD-ORFs were simultaneously amplified from 3μl yeast lysate, using well-specific primers (see table below). After PCR amplification, barcoded PCR products from an entire 96 well plate were pooled together and purified and sequenced with Illumina Solexa technology allowing for identification of interacting first pass pairs of proteins (FiPPs). To identify likely true AD/DB pairs, we developed a “SWIM score”^4^ *S* that takes into account the AD and DB reads in each well, total reads returned from the sequencing run, and other factors.

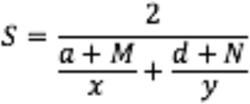

where *x* and *y* are read counts of an AD-ORF and DB-ORF in a given well respectively, *a* and *d* are total read counts of all aligned AD-ORF and DB-ORF in that well, and *M* and *N* are pseudo-counts for AD and DB respectively, which were constant for each sequencing batch but varied for different batches. We selected FiPPs for pairwise testing using a cutoff that balances the risk of testing too many false positives FiPPs versus not testing too many true positive FiPPs. The cutoff varied for different screens and sequencing runs to adjust for slight variations in the screening and sequencing protocol. Primers used for SWIM-seq are shown in **Suppl. Table 13**.

### Pairwise test of FiPPs

Each FiPP was subjected to a pairwise retest ^4^. Briefly, ADs and DBs were picked from the yeast ORF expression collection, mated in individual quadruplicates, and diploid yeast were spotted on selection media: SC-Leu-Trp-His+3AT and SC-Leu-His+3AT+CHX. Positive pairs were picked into SC-Leu-Trp and a SWIM-PCR was performed on the diploid yeast lysates to confirm the identity of ORFs. We used computational analysis as described in ^4^ to generate a list of binary interactors identified in the screens. If a DB acted as a *de novo* auto-activator, it was retested in a final pairwise experiment, where in parallel to mating the protein pairs, each DB was also mated against an “AD-null” plasmid without any ORF in the cloning site. Genes corresponding to mated yeast that grew on selective media when mated against AD-null yeast were removed from the final FlyBi dataset.

### Computation-based prediction of positive pairs and quality analysis

Computational prediction of positive pairs, based on the assay version 1 results (i.e., screens 1 and 2) was performed as described in ^37^. We used network-based link prediction to rank candidate interacting pairs based on the normalized number of length three network paths linking them (L3). As the input, we used a list of 2,195 PPIs from screens 1 and 2 and obtained the top 10,000 predictions. To quality-analyze these predictions and screen 2 data, we experimentally tested the top 1,000 predictions from the L3 computational analysis, a set of 135 positive interactions from screen 2 (positive benchmarks), 263 proteins from the RRS (negative benchmarks), and binary interaction pairs from the following sources or lists: Lit-BM-16 (see main text), and DPiM^41^. Altogether, we pairwise tested 3,399 non-redundant pairs in two orientations, allowing us to classify each pair as either positive, negative, or undetermined, following the experimental protocol described above and outlined in **Suppl. Fig. 1B**.

### MAPPIT validation

MAPPIT analysis was performed as previously reported^61^. Entry clones for bait and prey proteins were first cloned into MAPPIT vectors via Gateway LR reaction. Miniprep DNA was used to transfect HEK293T cells by standard calcium precipitation in quadruplicate. For each tested pair, two wells were left untreated and two were stimulated with the cytokine erythropoietin (Epo), which can induce JAK-STAT pathway signaling in cells in which there is a bait-prey interaction, resulting in activation of a STAT-responsive firefly luciferase reporter^61^. MAPPIT validation assays were only deemed valid if both bait and prey were successfully cloned into expression vectors and bait expression was detected. Fold-induction values (i.e., the signal from stimulated cells / signal from unstimulated cells) were calculated for each pair and two negative controls (i.e., no bait with prey and bait with no prey). Each tested pair was assigned a quantitative score comprising the fold-induction value of the pair divided by the maximum fold-induction value of the two negative controls. The validation was done in several batches, and the same ∼200 pairs of Lit-BM-16 (positive controls) and ∼200 RRS pairs (random controls) were included in each batch. Altogether, we validated 963 screen pairs, as well as 193 CuraGen high confidence pairs, 187 CuraGen low confidence pairs^2^, 216 pairs reported in^41^, 291 pairs in the “Finley Yeast Two-Hybrid Data” list that can be downloaded from the DroID online resource^27^, and 188 Lit-BS pairs.

Pairs were scored positive or negative based on thresholds set at the 99th percentile of the RRS scores (equivalent to a 1% false discovery rate). Each experimental batch was scored separately and used the quantile function in the Python library. Pairs without valid quantitative scores were dropped, and recovery rates were calculated as the number of positive pairs over the sum of the positive and negative pairs. The error bars on the recovery rates were standard errors of the portions.

### Bioinformatic analyses

#### SAFE analysis

For SAFE analysis, we used the SAFE software^62^ v1.5 to determine and visualize significant functional modules in various networks. The network layouts were generated with Cytoscape^63^ v3.4.0 using the edge-weighted spring embedded layout. SAFE analysis was run using the default options with the exception that “layoutAlgorithm” was set to “none” (using the layout as generated by Cytoscape) and the “neighborhoodRadius” was set to “2.”

#### Gene set enrichment

Gene set enrichment analysis of genes covered by the FlyBi network was done using an in-house program written based on a hypergeometric distribution test. Gene sets were built based on the Gene List Annotation for *Drosophila* (GLAD) database^64^. A negative control of 1000 random networks was generated by shuffling FlyBi gene nodes 1000 times.

#### Identification of Lit-BM-20

Lit-BM-20 was built by selecting *Drosophila* physical interactions from MIST for which either the interaction was identified using one detection method for direct physical interaction as reported in multiple publications or the interaction was identified using multiple methods for detection of direct interactions (or both). The list of the detection methods for direct interaction were annotated based on the same criteria used for building the HuRI network^4,38^. Annotations of detection methods for interactions included in MIST were based on the European Bioinformatics Institute (EBI) molecular interaction (MI) controlled vocabulary system (https://www.ebi.ac.uk/ols/ontologies/mi), for example, MI:0800 for two hybrid.

#### Comparisons to interologs

The FlyBi dataset was compared with PPIs and genetic interactions detected in *Drosophila* and interologs as assembled by MIST. In addition, the FlyBi dataset was compared with *Drosophila* orthologous gene pairs mapped using DIOPT ^65^ from yeast gene pairs with similar genetic interactors^66^.

#### Adjacency matrix

An adjacency matrix for binary interactions was built using FlyBi interactions and Lit-BM-20 interactions to visualize how frequently interacting proteins are reported in literature. The interacting proteins were binned and ordered along both axes based on the number of corresponding publications. The color intensity of each square reflects the total number of interactions between proteins in the corresponding bins.

#### Gene Ontology (GO) analysis

Analysis of the biological relevance for interacting proteins was done by evaluating commonality in the GO annotation, phenotype annotation, and complex memberships of the interacting proteins from FlyBi and Lit-BM-20 data as compared with protein pairs from the random networks. To do this, gene2go and gene2phenotype annotations were obtained from FlyBase^67^, and GO terms with more than 30 associated genes were removed prior to enrichment analysis.

#### Protein complex-based analysis

Complex-based interaction data for *Drosophila* were obtained from MIST^29^. Protein complex annotations were obtained from COMPLEAT^42^. COMPLEAT includes annotated complexes from the literature and complexes predicted based on the connectivity of protein-protein network; however, only literature-based complexes were used for enrichment analysis.

#### Co-localization analysis

Co-localization analysis was done based on organelle prediction by PSORT^68, 69^ and DeepLoc^70^. Co-citation analysis was done based on associated literature for each interacting protein. Genome-scale studies were removed and only publications with fewer than 100 associated genes were considered. Co-expression analysis was done by mining a single-cell RNA-seq dataset for the *Drosophila* midgut^71^ to identify cell types in which each interacting protein is expressed. The results were visualized by plotting the fraction of interacting pairs that share the same organelle annotation, the number of interacting pairs cited in the same publication(s), or the average number of co-expressed cell types of the interacting pairs, in each case as compared to results with the 1,000 randomized networks.

### Autophagy-related assays

#### Drosophila strains and genetic assays

Flies were raised at 25°C following standard procedures unless otherwise noted. The following *Drosophila* strains were used: *UAS-Atg1* (BDSC51655), *dwg^8^* (BDSC4094), *r4-mCherry-Atg8a Act>CD2>GAL4 UAS-GFP-nls*^58^, *UAS-Luc-RNAi* (BDSC31603). Additional *Drosophila* strains used for the genetic screen are listed in **Suppl. Table 10**.

#### In vivo autophagy assay in adults and comparison of autophagy and random sets

To compare datasets of comparable size, 106 nodes from a randomly generated network were selected as a control gene set and one stock per gene was screened for the modifier of *Atg1* overexpression-induced eye phenotypes. To compare the autophagy set covered by multiple stocks per gene with the random set where only one stock per gene was used, we randomly selected one stock per gene for the autophagy set and compared it with the result from the random set. We repeated this comparison process five independent times.

#### In vivo autophagy assay in larvae

Second instar larvae were collected 72-96 hr after egg laying and cultured in fresh fly media with yeast paste (fed) or in vials containing 20% sucrose (starved) for 4 hrs. Autophagy level is indicated by autophagosome numbers labeled by mCherry-ATG8a. GFP-marked clones expressing RNAi or protein in the larval fat body were generated through heat shock-independent induction as previously described^52^.

#### Immunofluorescence assays

Dissected fat bodies were fixed in a solution of 4% PFA/PBS for 40 minutes. After permeabilization with 0.3% Triton/PBS, fat bodies were washed, and incubated overnight with anti-HA antibodies, and visualized using anti-mouse Alexa-633 (Invitrogen). S2R+ cells expressing Flag-dwg were fixed with 4% paraformaldehyde, permeabilized with 0.1% triton, and processed for immunostaining. DAPI (1 μg/ml) was used to stain nuclei. Samples were examined using a Zeiss LSM 780 confocal laser scanning microscope (Carl Zeiss Inc.) with a 63x Plan-Apochromat (NA1.4) objective lens.

### Co-Immunoprecipitation analysis in *Drosophila* cells

#### Plasmids

Full-length ORFs of *CG11486* (GEO01712), *CG9667* (GEO12785), *Deaf1* (GEO12259), *RfC4* (GEO04321), *CG7006* (GEO04456), *Larp* (GEO13890), *CG4813* (GEO05615), *lwr* (GEO03784), *DOR* (GEO05909), *dwg* (GEO06061), *wash* (GEO08420), *mri* (GEO13088), *me31B* (GEO01853), *MED15* (GEO05444), *Nup54* (GEO04647), *sm* (GEO09592), *smB* (GEO05072), *CG10209* (GEO04957), *MED4* (GEO04489), *Odj* (GEO06447), *Dpy-30L2 (GEO12106), MED19 (GEO06036), Chro* (GEO02531), *Atg8b* (GEO01803), *Atg8a* (GEO03266), *CG5446* (GEO09660), *CG4813* (GEO05615), *kin17* (GEO12682), *CG7484* (GEO09148), *sala* (GEO07724), and *CG9667* (GEO12785), from the FlyBi ORF clone collection reported in this work. ORFs were transferred into the *Drosophila* gateway vectors pAWF, pAGW or pAWG. The GFP ORF was cloned into pAWM as a control. To generate the Dwg deletion mutant proteins, DNA sequences corresponding to amino acids 1-150, 134-215, 215-393, and 385-592 of *dwg* were PCR amplified and subcloned into the pAWF vector. Using PCR mutagenesis, we generated *dwg^Y129A-I132A^, dwg^F401A-L404A^,* and *dwg^Y129A-I132A-F401A-L404A^* mutants by replacing tyrosine 129, isoleucine 132, phenylalanine 401, or leucine 404 with alanine, followed by cloning into pAWF or pTWH. Mutant ORF sequences were verified by Sanger DNA sequencing.

#### Antibodies

Antibodies used for the study were as follows: anti-GFP (Molecular Probes, A6455), anti-Atg8 (Abcam, ab109364), anti-Flag (Sigma, F3165), and anti-GAPDH (GeneTex, GTX100118).

#### Cell culture

*Drosophila* cells were cultured in Schneider’s medium supplemented with 10% fetal bovine serum (FBS) at 25°C. For Rapamycin (LC Laboratories, R-5000) or Bafilomycin (Sigma, B1793) treatment, S2R+ cells were treated with 20 nM Rapamycin or 100 nM Bafilomycin-A1 (Baf-A1) for 24 hrs.

#### Immunoprecipitation and immunoblotting

DNA was transfected into S2R+ cells in a 10cm plate with Effectene transfection reagent (Qiagen) following manufacturer’s protocol. After 3 days of incubation, cells were lysed using lysis buffer (Pierce) with protease inhibitor (Thermo Fisher Scientific) and phosphatase inhibitor (Sigma). Lysate was incubated with GFP-Trap agarose beads (Bulldog Bio) or anti-Flag M2 magnetic beads (Sigma) for 2 hr at 4°C to precipitate the protein complexes. Beads were washed 3-4 times with 1 ml lysis buffer. SDS-sample buffer was added, and the samples were boiled at 95 °C for 10 min. Boiled samples were run on polyacrylamide gel (Bio-Rad) and transferred to Immobilon-P polyvinylidene fluoride (PVDF) membrane (Millipore). The blot was probed with primary antibody, followed by HRP-conjugated secondary antibody, and signal was detected by enhanced chemiluminescence (ECL; Amersham).

#### Quantification of mRNA expression

Total RNA was extracted from control or *dwg^8^* mutants using TRIzol® reagent (Invitrogen). We synthesized the first strand cDNA with 1 µg of total RNA using iScriptTM Reverse Transcription Supermix (BIO-RAD) followed by quantitative PCR with CFX96 Real-Time System (BIO-RAD) using iQTM SYBR Green Supermix (BIO-RAD). All expression values were normalized to *RpL32* (also known as *rp49*). All assays were performed in triplicate. The primer sequences used for PCR are as follows: Rp49: ATCGGTTACGGATCGAACAA, GACAATCTCCTTGCGCTTCT Atg1: CTAAAGCCGTCGTCCAATGT, GAACAGCATGCTCCGGTATT Atg17: GAAGCTGCACAACATCCCG, GCTGAGTAGCACGACACTTGG Atg3: CCAAGACAAAACCCTACCTACC, GCCGACGTATTCCATCTGCT Atg13: GAACCTAAAGACAGGAGAGAGCA, ACCCTCAGTCGTTTTCAGGGA

### Chromatin immunoprecipitation (ChIP)

S2R+ cells expressing Flag-dwg were subjected to ChIP assays using SimpleChIP Plus Enzymatic Chromatin IP Kit (Cell Signaling Technology) according to the manufacturer’s protocol. DNA co-immunoprecipitation with either anti-Flag antibody or IgG control antibody was analyzed by deep DNA-sequencing or quantified by ChIP-qPCR using primers shown below.

Atg1: CACTTGCAGGATCGATGGCA, TTACGCTGATCGTCCGTGTG

Atg17 promoter: CACATGCTCGGCCTGCTATT, CAGACTGTCGCTGGTGCTTT

Atg17 intron: TGCCCGCATCGTGTAAATGG, CTGCTGCTGCTGTGAGTGTT

Atg3: AGCTGCGAAGTGCAAGTCAA, GCGTCAGATATTCGGCCACA

Atg13: AATCGCAGTGAAAGGGCGTT, AGTTCGCTGTCTGCGTTTGT

For ChIP-seq data analysis, low-quality reads and adaptor primer sequences were trimmed using Trim Galore 0.6.4 (https://github.com/FelixKrueger/TrimGalore), and trimmed reads were mapped against fly genome dm6 by bowtie2 2.3.5.1 with the additional argument “-q --local”^72^. Samtools 1.6 were used to sort, filter unique reads, and convert file format to bam files^73^. Peak calling was performed with MACS2 2.2.6 using the additional parameter “-B --SPMR -f BAMPE -g dm”^74^. Peaks were annotated with HOMER 4.11^75^. DeepTools 3.4.0 were used for normalizing read counts to CPM and convert bam files to bigWig format^76^.

### Plasmid, data, and code availability

#### Plasmid clones and sequence data

Plasmid clones are available from both the *Drosophila* Genomics Resource Center (University of Indiana, Bloomington, IN) and the DNASU plasmid repository (Arizona State University, Phoenix, AZ). The ORF in the Gateway donor vector were end-read sequenced (see “Generation of a large-scale ORF clone resource”). For a subset of 954 ORFs, the end-reads sequence spanned the full ORF. For this subset, we submitted the sequence data to the NCBI GEO database.

#### FlyBi data

FlyBi binary interaction data are available downloadable data file at the FlyBi project webpage <http://flybi.hms.harvard.edu/>. In addition, these pairs have been integrated with other datasets at IntAct <https://www.ebi.ac.uk/intact/>39 and in the Molecular Interaction Search Tool (MIST) <https://fgrtools.hms.harvard.edu/MIST/>29. MAPPIT data as well as interaction and RNAi data for the autophagy-related network are available in **Suppl. Tables 6, 7, 8** and **10.**

#### Code availability

The L3 prediction code, together with example datasets, input data files and predictions, is available at <https://doi.org/10.5281/zenodo.2008592>.

## Figures & Figure Legends

**Supplemental Figure 1:**
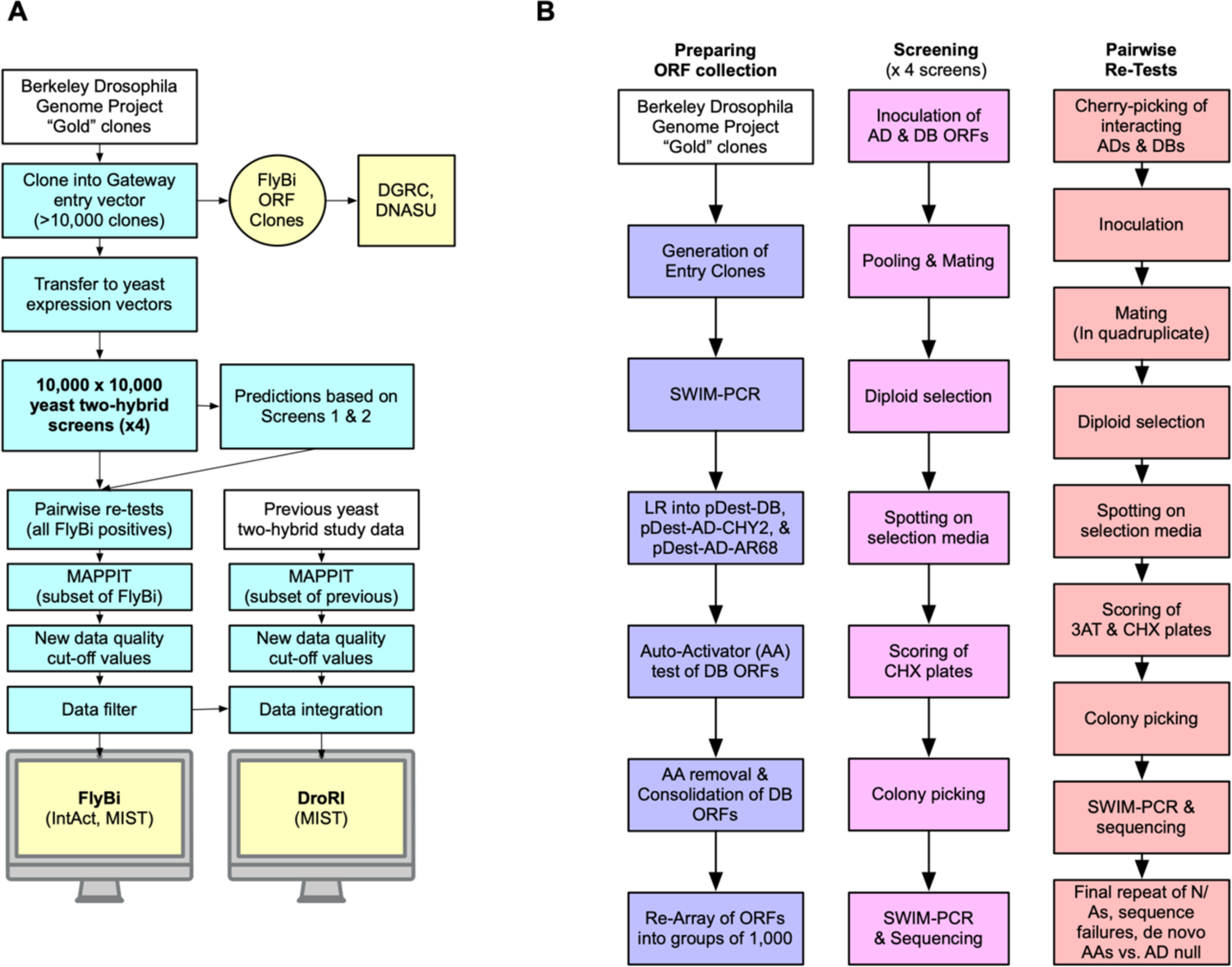
FlyBi project workflow and experimental yeast two-hybrid (Y2H) approach. **A.** Overall project workflow. Pre-existing physical resources or datasets are shown in white. New experiments or analyses are shown in blue. Newly generated physical or data resources, or their locations, are shown in yellow. The project resulted in a new experimentally determined Y2H dataset (FlyBi) and a new *Drosophila* reference interactome (DroRI). **B.** Detailed Y2H screen workflow. Left, generation of the ORF collection. ORFs were amplified with M13 SWIM primers and successfully sequenced ORFs were transferred into yeast expression vectors via Gateway cloning and transformed into yeast (pDEST-DB into Y8930, pDEST-AD-*CYH2* and pDEST-AD-AR68 into Y8800). ORFs in pDEST-DB were tested for auto-activation by spotting on Synthetic Complete media lacking Leucine and Histidine (SC-Leu-His) plates. Strong auto-activators (AA) were removed and the yeast ORFs were re-arrayed into groups of 1,000. Center, screening pipeline. AD and DB ORFs were inoculated in corresponding selection media and incubated at 30°C overnight. The next day, 1,000 AD-ORFs were pooled to obtain a kilo-pool and mated against a single DB in YPD (yeast extract peptone dextrose) media. After overnight incubation at 30°C, 10 µl of the mated yeast culture was transferred into Synthetic Complete media lacking Leucine and Tryptophan (SC-Leu-Trp media) to select for diploid yeast cells. The following day, diploid yeast cells were spotted on SC-Leu-Trp-His+1mM 3AT selection media and SC-Leu-His+1mM 3AT+CHX. Colonies growing on SC-Leu-Trp-His+1mM 3AT plates but not on SC-Leu-His+1mM 3AT+CHX plates, were picked up to 3 times from each spot. Lysates, SWIM PCR and sequencing were performed to identify interacting pairs. Right, pairwise testing. Interacting pairs were arrayed into 96-well plates for AD and DB respectively, inoculated into selection media and after incubation overnight at 30°C mated in quadruplicates. The next day, diploid selection was performed by transferring 10 µl mated yeast culture into SC-Leu-Trp. After growth overnight at 30°C, diploid yeast cells were spotted on selection media. For positive interactions growing on 3AT plates only and not on selection plates containing CHX, one out of four replicates was picked, SWIM PCR performed and sequence confirmed. A final pairwise test was performed for pairs that scored as de novo auto-activators or NAs (not applicable due to invalid scores) in the pairwise test. For those pairs, each DB-ORF was separately mated with an “AD-null” plasmid (no ORF in the cloning site) in parallel with the AD interactor. If a yeast colony grew more strongly when mated to the corresponding AD-ORF than the AD-null strain, the PPI was considered valid and added to the dataset after sequence confirmation. Abbreviations: SWIM PCR Shared-Well Interaction Mapping by sequencing; AA Auto-Activator; SC-Leu-Trp-His+1mM 3AT Synthetic Complete-Leucine-Tryptophan-Histidine+ 3-amino-1,2,4-triazole; SC-Leu-His+1mM 3AT+CHX Synthetic Complete-Leucine-Histidine+ 3-amino-1,2,4-triazole+ Cycloheximide; SC-Leu-Trp, Synthetic Complete-Leucine-Tryptophan.

**Supplemental Figure 2:**
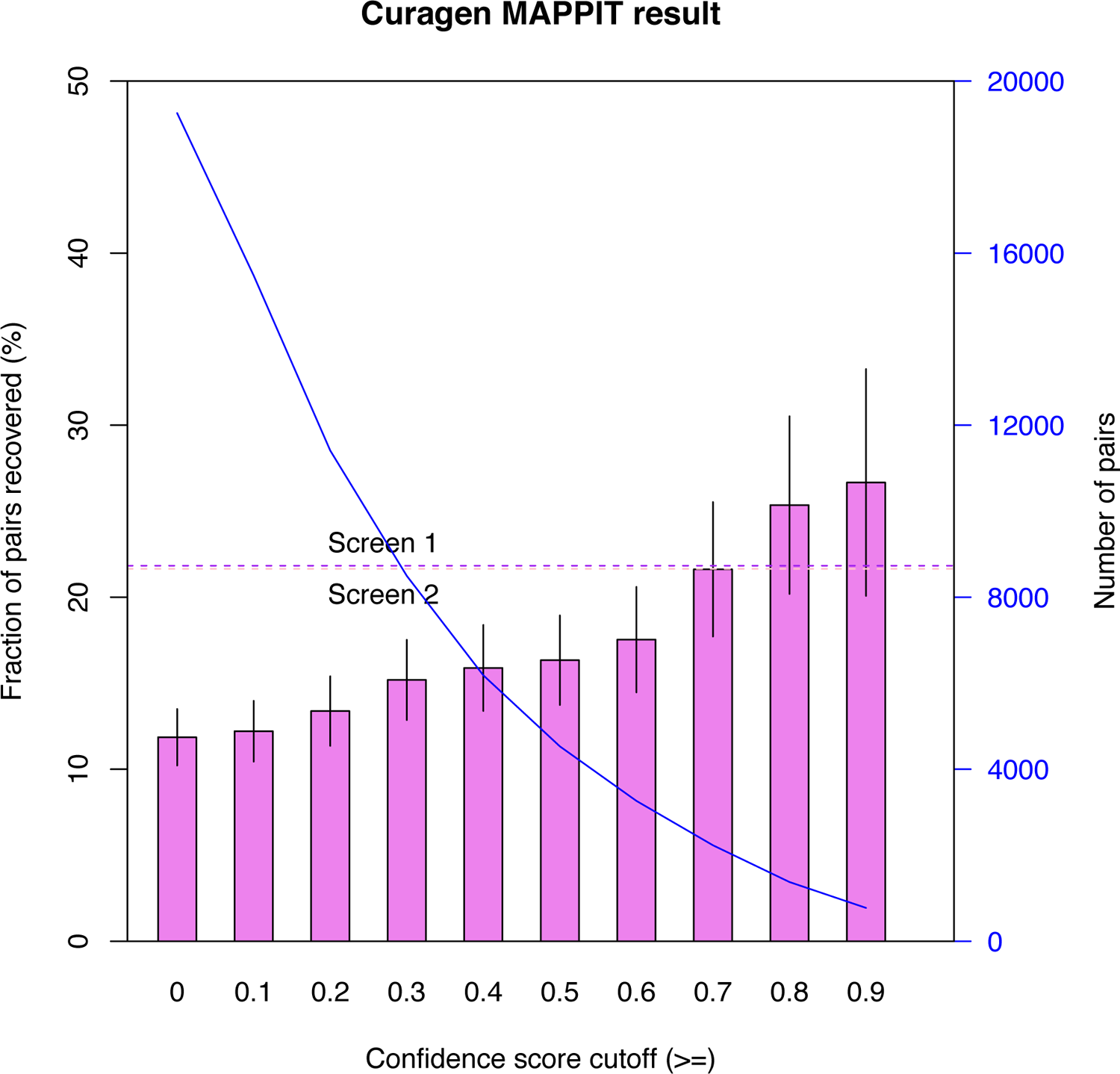
Experiment evaluation of CuraGen pairs at different CuraGen confidence score cutoffs in MAPPIT assay. Blue curve, the declining number of pairs left in the dataset at the indicated CuraGen stringency cutoffs are applied across all pairs. Dashed line, the recovery rates of FlyBi screens 1 and 2 (i.e., the FlyBi screens performed in the most comparable screen version).

**Supplemental Figure 3:**
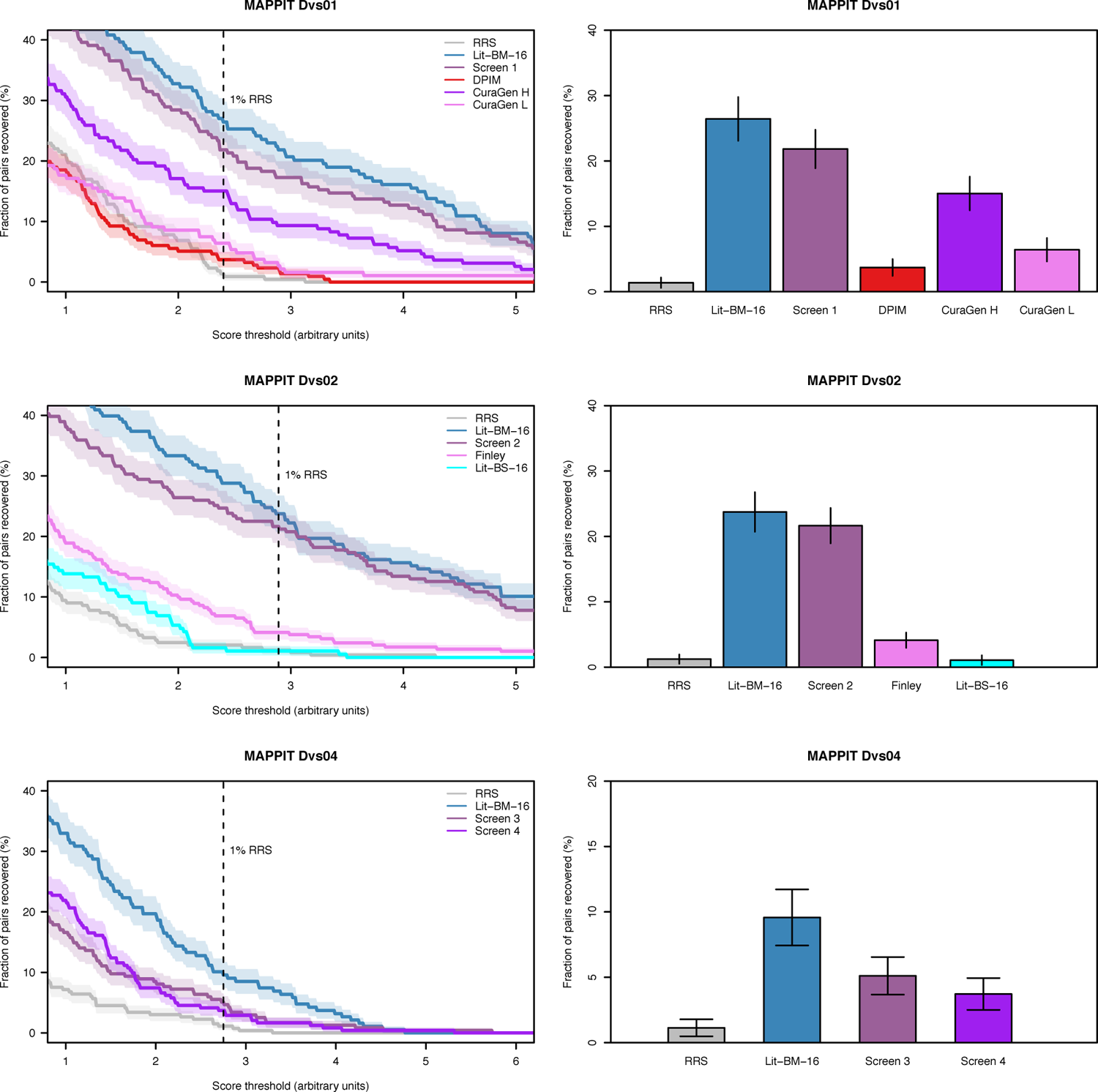
Experimental validation of pairs from FlyBi and other datasets in the MAPPIT assay. The MAPPIT configurations were N-N (top and middle) or N-C (bottom), in accordance with configurations of version 1 (N-N) and version 3 (N-C) in the FlyBi Y2H screens. Clouds around the solid lines indicate the standard of error. Left panels, titration plots showing the recovery rates at different thresholds. Right panels, bar plots showing the recovery rate at a cutoff value for which the random reference set (RRS) scores at a rate of 1%. Error bars indicate the standard of error.

**Supplemental Figure 4:**
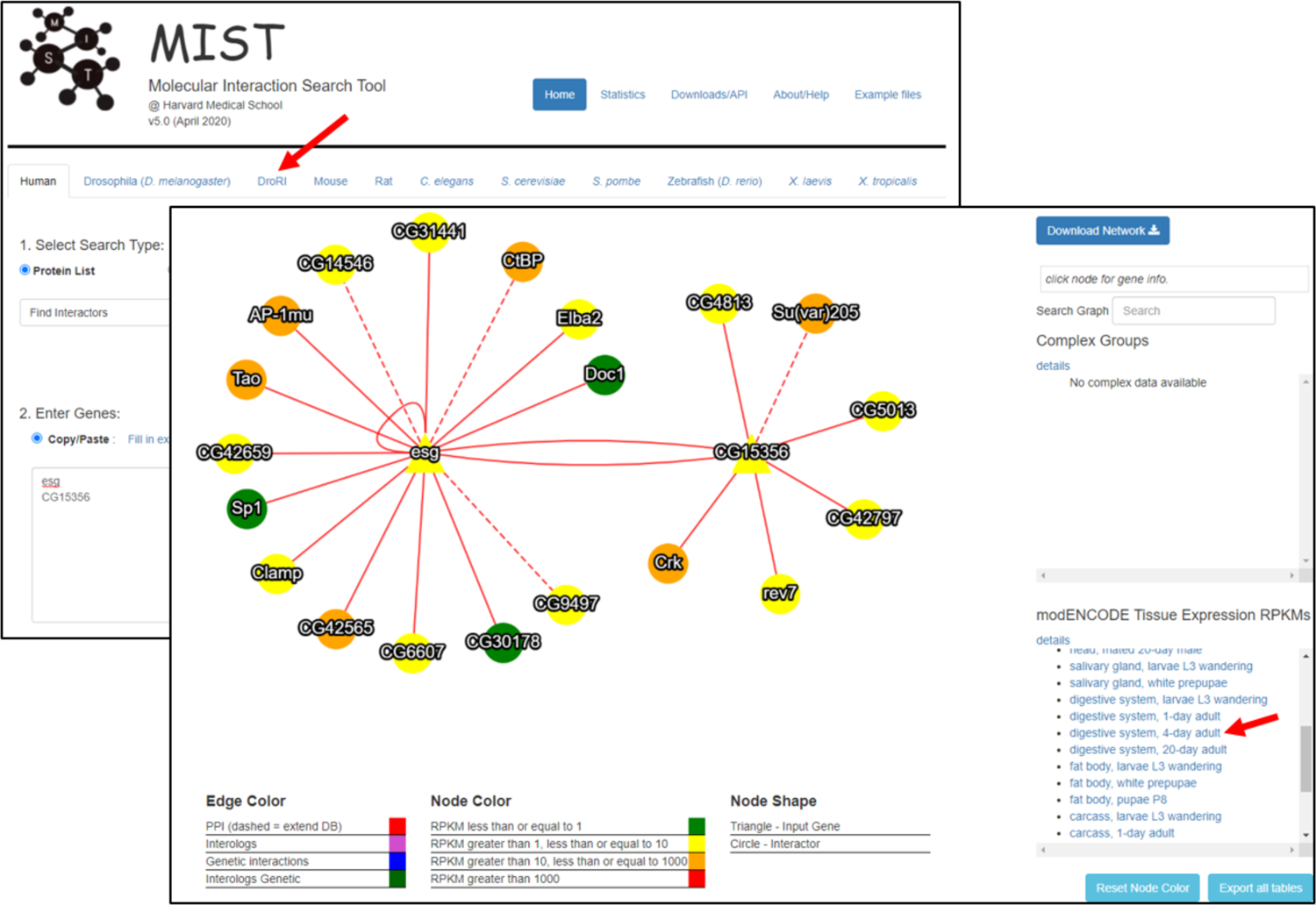
Integration of DroRI network and transcriptomics data at MIST. Interactions in the DroRI master network can be queried from a dedicated tab at MIST (top left, red arrow). Users have the option to project expression levels as reported by modENCODE tissue-specific RNA-seq datasets onto the network (bottom right, red arrow). The search input is either a gene or a list of genes. An example results page for a query with *escargot* (*esg*) and *CG15356* is shown. Solid edges, binary interactions identified in the FlyBi screens; dotted edge, binary interactions curated from the literature. Node colors reflect expression levels for the selected dataset. In this example, the network is overlayed with modENCODE RNA-seq data for the digestive system of 4-day adults.

**Supplemental Figure 5:**
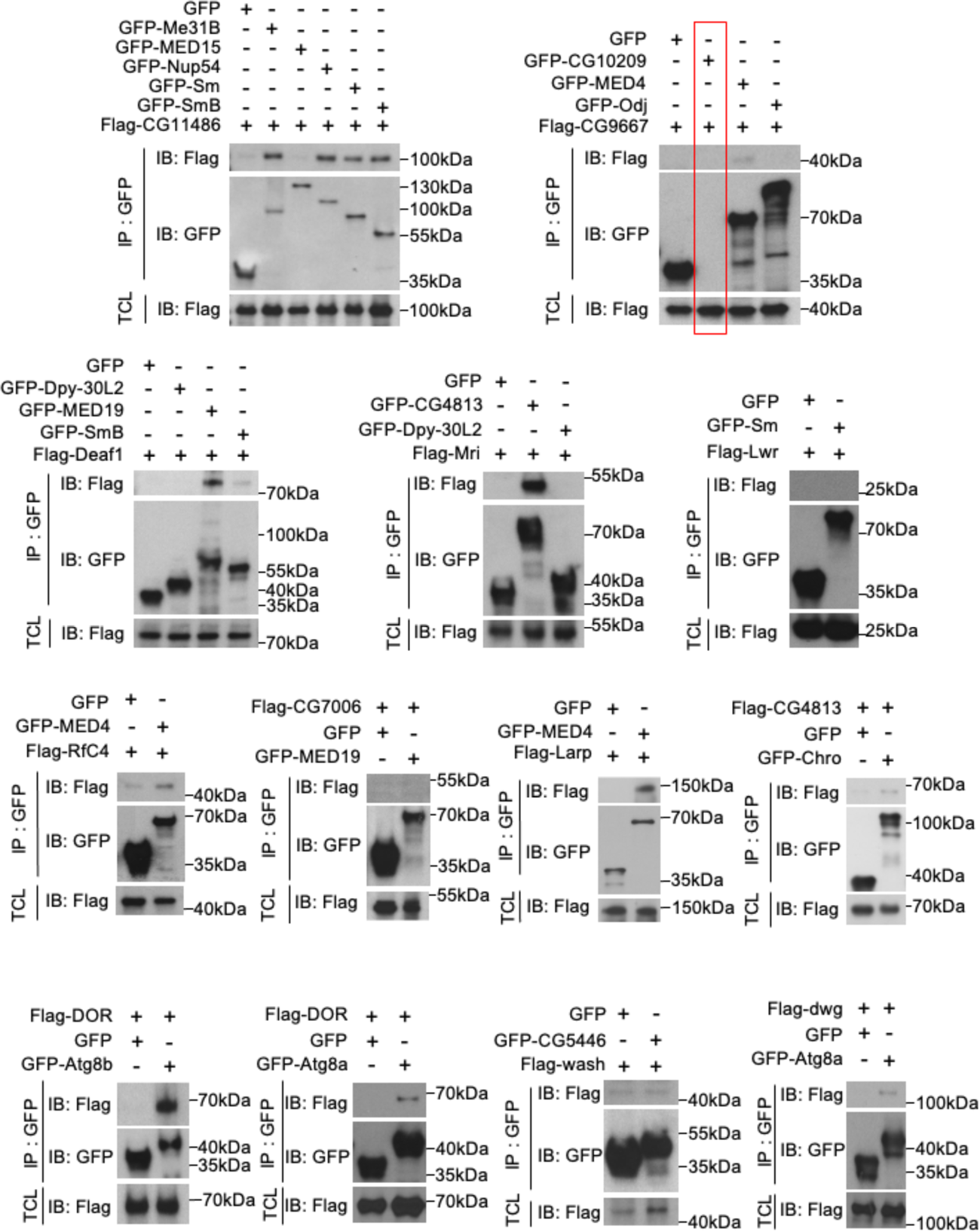
Experimental validation of interactions between putative autophagy-related proteins by co-immunoprecipitation. Open reading frames (ORFs) were fused in-frame with epitope tags or GFP, co-expressed in *Drosophila* cells, and subjected to co-IP followed by detection with antibodies as indicated. See also summary of results in **Suppl. Table 11** and network visualization in **Fig. 4B**.

**Supplemental Figure 6:**
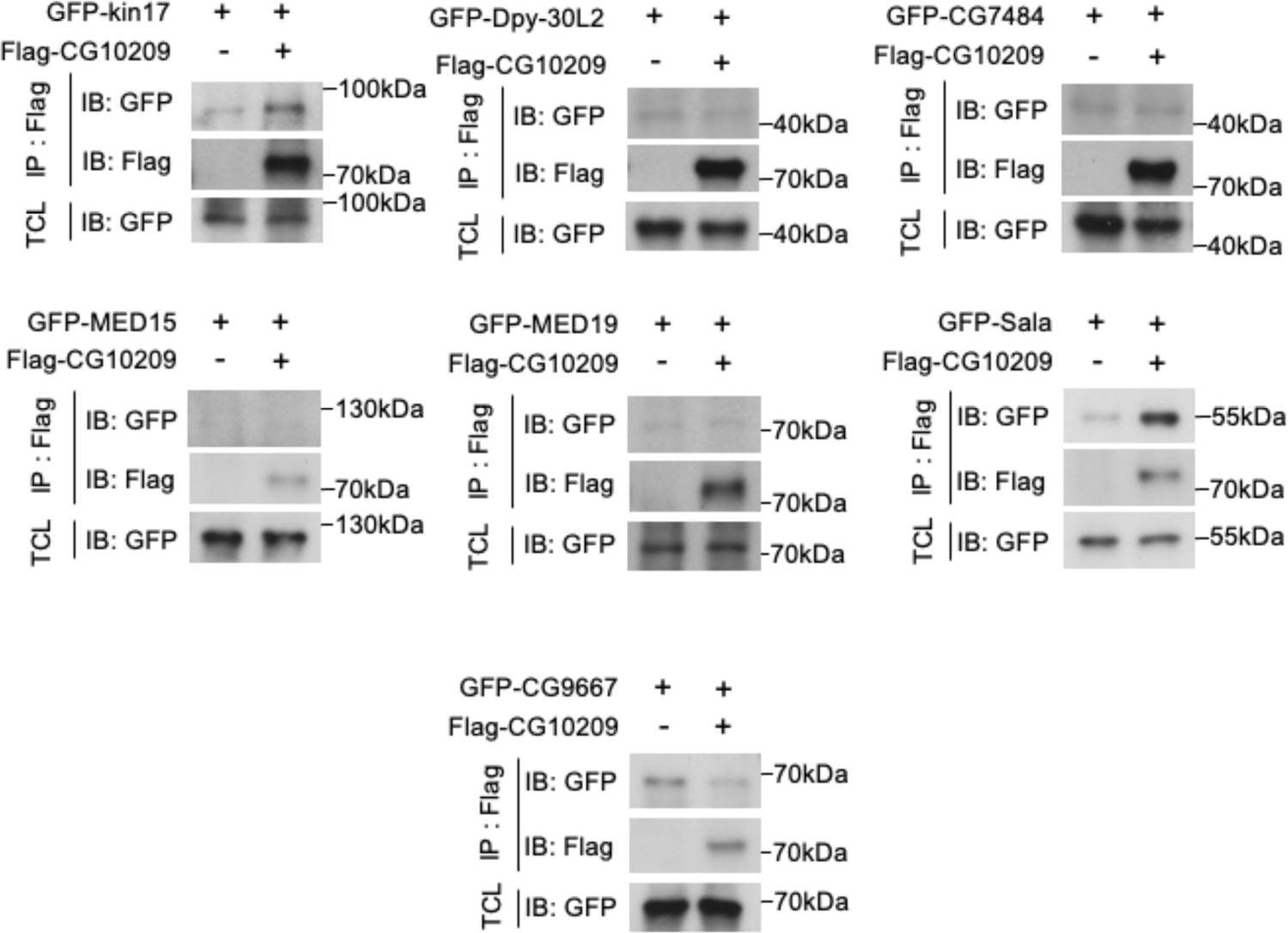
Experimental validation of interactions between CG10209 and other putative autophagy-related proteins by co-immunoprecipitation (co-IP). As GFP-tagged CG10209 is undetectable (Supplemental Figure COIP-1), we designed a smaller Flag-tagged form of CG10209 and enriched it using Flag-beads for immunoprecipitation. Open reading frames (ORFs) were fused in-frame with epitope tags or GFP, co-expressed in *Drosophila* cells, and subjected to co-IP followed by detection with antibodies as indicated. See also summary of results in **Suppl. Table 11** and network visualization in **Fig. 4B**.

**Supplemental Figure 7:**
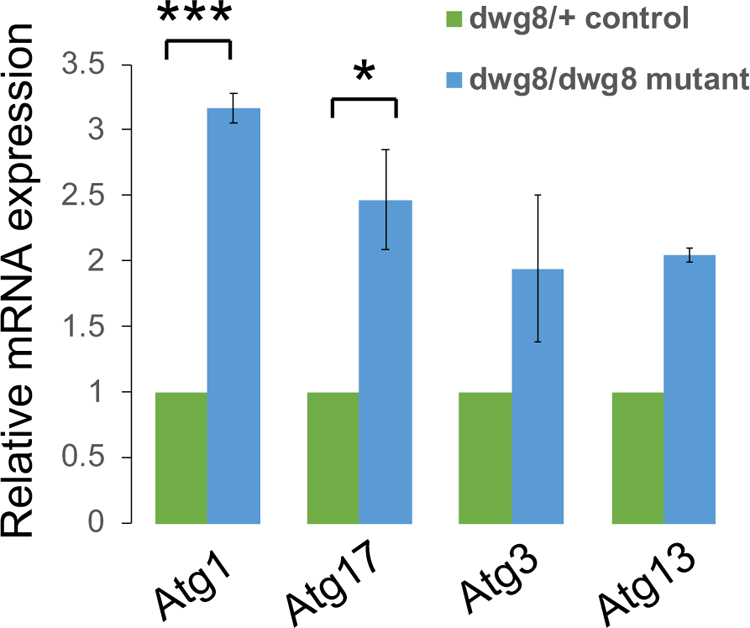
Increased expression of *Atg* transcripts in *dwg* mutants. Relative mRNA expression of *Atg1, Atg17, Atg3*, and *Atg13* genes in control (*dwg^8^/+*) and *dwg* mutants (*dwg^8^/dwg^8^*). Measurements shown are mean ± SEM. One-Way ANOVA followed by Tukey’s multiple comparisons test was performed to identify significant differences; Data is expressed as means ± SEM of three independent experiments; ***P<0.001, *P<0.05.

**Supplemental Figure 8:**
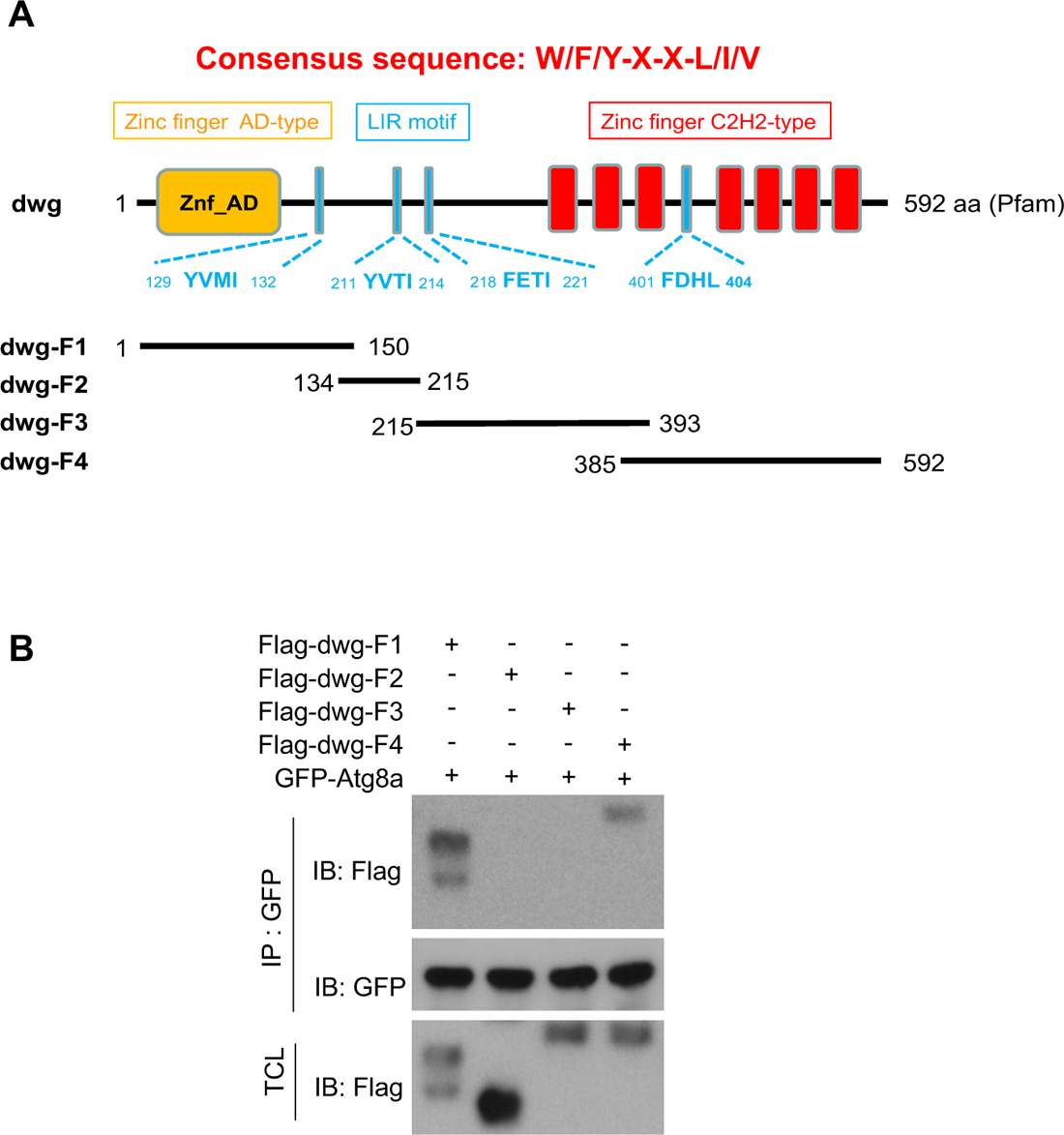
Mapping the Dwg-Atg8a interaction regions. (A) Schematic representation of the domain structures and LIR (LC3-interacting region) motifs, and deletion mutants of *dwg*. (B) Atg8a interacts with the first and fourth LIR motifs of dwg. S2R+ cells transfected with *GFP-Atg8a* and *Flag-dwg-F1-4* for 48 hr followed by immunoprecipitation with anti-GFP nanobody. The immunoprecipitated proteins and total cell lysates were analyzed by immunoblotting with antibodies as indicated.

**Supplemental Figure 9:**
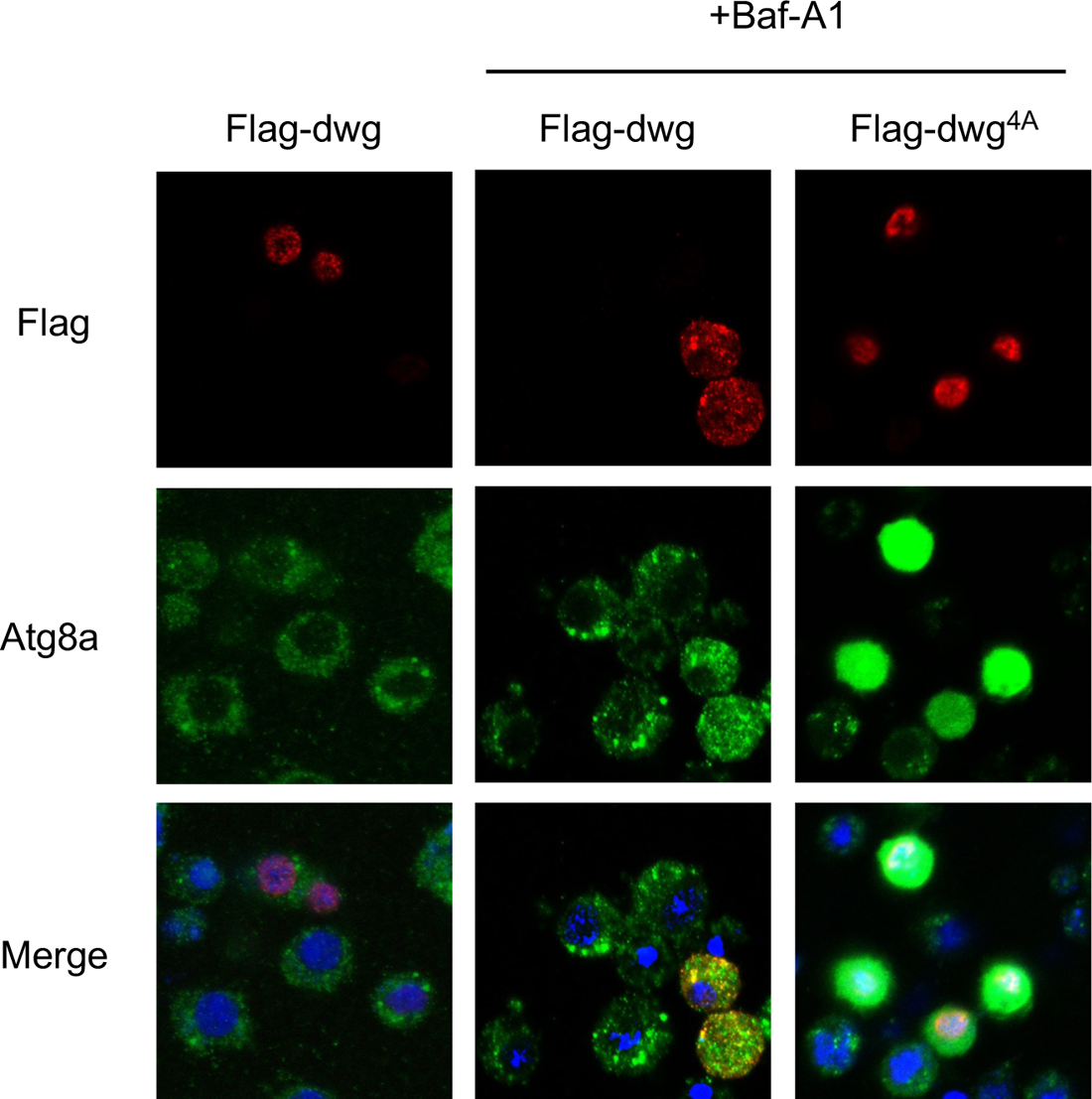
Disruption of the interaction between Dwg and Atg8a inhibits autophagy. In untreated S2R+ cells, Dwg is localized to the nucleus. Bafilomycin A1 (Baf-A1) treatment inhibits autophagosome degradation and increases detectable Dwg in the cytoplasm and co-localization of Dwg with Atg8a punctae. A Dwg variant with LIR motif mutations (Dwg^4A^) is restricted to the nucleus and strongly suppresses autophagy.

**Suppl. Fig. 10:**
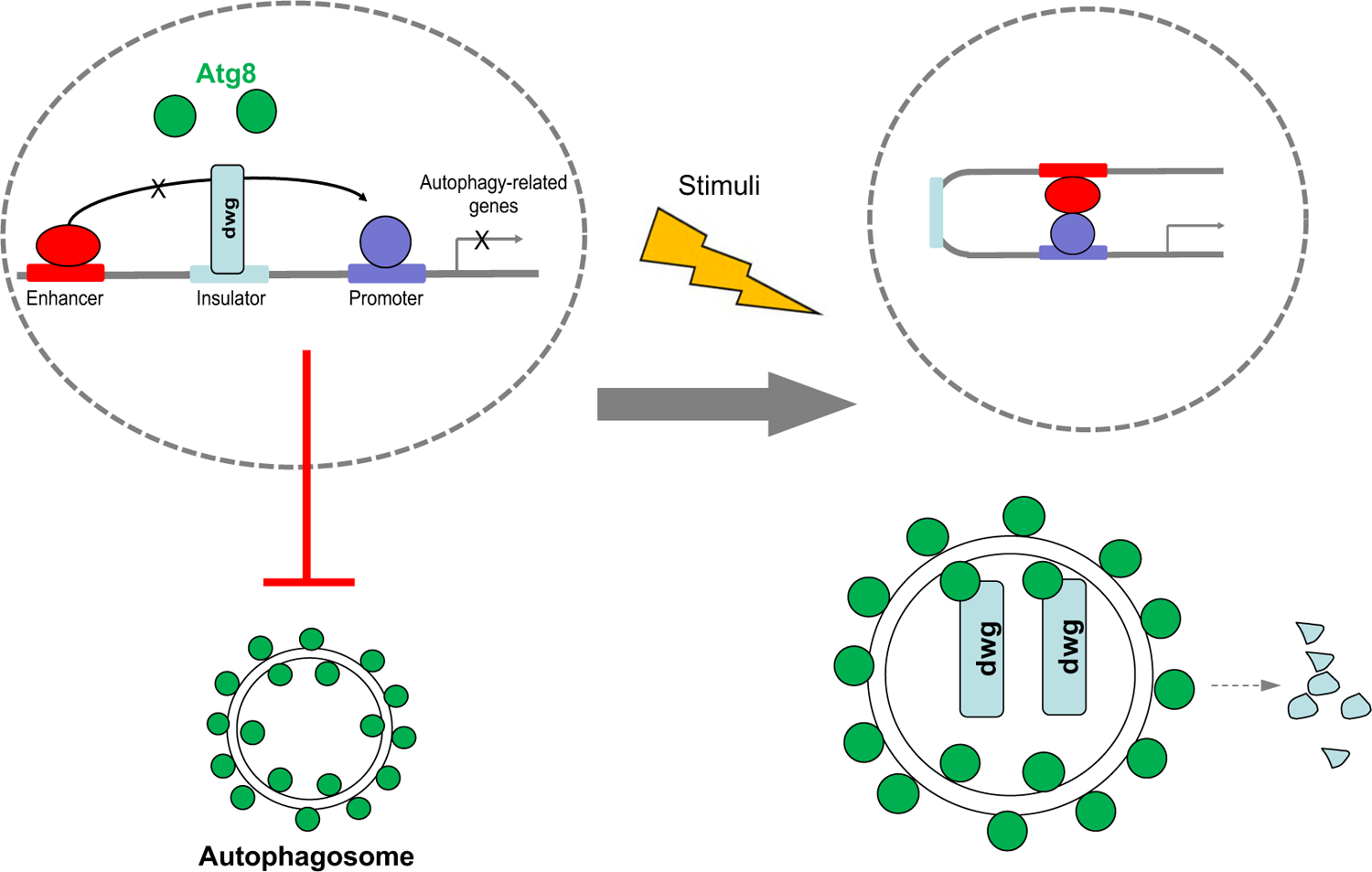
Working model of regulation of Dwg by Atg8a/autophagy. Under normal conditions, Dwg binds to insulator elements and inhibits transcription of *ATG* genes. Following a stimulus such as starvation, autophagy is induced, and Atg8a associates with and transports Dwg from the nucleus to autophagosomes for degradation.

## Supplemental Tables

**Suppl. Tables 1-10 are included as separate files**

**Supplemental Table 11:** Results of co-immunoprecipitation in *Drosophila* cells of Flag- and GFP-tagged proteins from the putative autophagy network.

| Flag-tagged protein (size<sup>1</sup>) | GFP-tagged protein (size<sup>1</sup>) | Interaction Detected by CoIP |
| --- | --- | --- |
| CG11486 (100) | Me31B (91) | yes |
| CG11486 | MED15 (120) | no |
| CG11486 | Nup54 (104) | yes |
| CG11486 | Sm (92) | yes |
| CG11486 | SmB (61) | yes |
| CG9667 (45) | CG10209 (96) | no |
| CG9667 | MED4 (68) | yes |
| CG9667 | Odj (90) | no |
| Deaf1 (76) | Dpy-30L2 (50) | no |
| Deaf1 | MED19 (75) | yes |
| Deaf1 | SmB (61) | yes |
| RfC4 (52) | MED4 (68) | yes |
| CG7006 (36) | MED19 (75) | no |
| Larp (121) | MED4 (68) | yes |
| CG4813 (59) | Chro (141) | yes |
| Lwr (33) | Sm (92) | no |
| DOR (60) | Atg8b (54) | yes |
| DOR | Atg8a (54) | yes |
| Dwg<sup>2</sup> (100) | Atg8a (54) | yes |
| Wash (68) | CG5446 (49) | no |
| Mri (63) | CG4813 (84) | yes |
| Mri | Dpy-30L2 (50) | no |
| CG10209 (75) | Kin17 (95) | yes |
| CG10209 | Dpy-30L2 (50) | no |
| CG10209 | CG7484 (60) | no |
| CG10209 | MED15 (120) | no |
| CG10209 | MED19 (75) | no |
| CG10209 | Sala (54) | yes |
| CG10209 | CG9667 (70) | no |

<sup>1</sup> Observed size of the fusion protein with tag, in kDa.
<sup>2</sup> Expected size, 81 kDa. The observed size of our Dwg fusion protein (100 kDa) is consistent with a report by <sup>55</sup>, who similarly observed that Dwg migrates at a higher molecular weight than expected.

**Supplemental Table 12:** Attributes of the yeast expression vectors used in this study.

| Plasmid Vector | pDEST-DB | pDEST-AD-CHY2 | pDEST-AD-AR68 |
| --- | --- | --- | --- |
| **Fusion** | Gal4-DB<br>(aa 1-147) | Gal4-AD<br>(aa 768-881) | Gal4-AD<br>(aa 768-881) |
| **Fusion location** | N-term | N-term | C-term |
| **Promoter** | Truncated *ADH1* promoter<br>(-701 to +1) | Truncated *ADH1* promoter (-701<br>to +1) | Truncated *ADH1* promoter (-410<br>to +1) |
| **Yeast replication origin** | CEN | CEN | 2 micron |
| **Linker** | SRSNQ | GGSNQ | VDGTA |
| **Terminator** | *ADH1* Term | *ADH1* Term | *ADH1* Term |
| **Selection marker** | AmpR | AmpR | AmpR |

DB, DNA binding domain; AD, activation domain; N-term, amino-terminal fusion; C-term, carboxy-terminal fusion; CEN, centromere; *ADH1* Term, *ADH1* terminator sequence; AmpR, ampicillin resistance.

**Suppl. Table 13:**
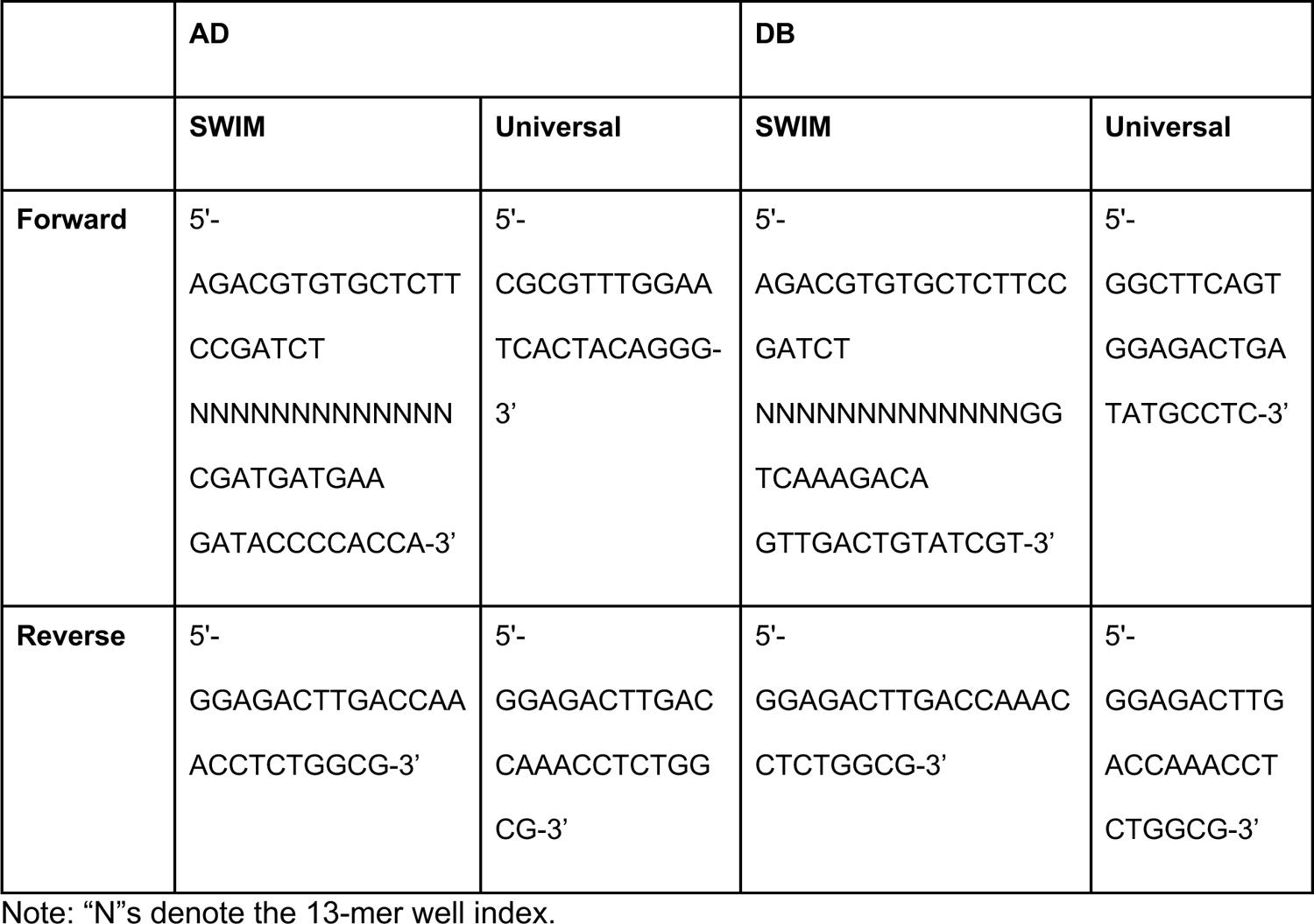
Primers used for SWIM-seq.

## Acknowledgments

We thank Jonathan Zirin for helpful comments. This work was supported by NIH NHGRI 5R01HG007118 and NIH NIAMS R01 AR057352 (N. Perrimon, PI). Additional support includes start-up funds from the Duke-NUS Medical School (R-913-200-171-263, to H.-W.T.) and a Foundation Grant from the Canadian Institutes of Health Research (to F.R.). NP is an investigator of the Howard Hughes Medical Institute. This article is subject to HHMI’s Open Access to Publications policy. HHMI lab heads have previously granted a nonexclusive CC BY 4.0 license to the public and a sublicensable license to HHMI in their research articles. Pursuant to those licenses, the author-accepted manuscript of this article can be made freely available under a CC BY 4.0 license immediately upon publication.

